# Altered metal distribution in the *sr45-1* Arabidopsis mutant causes developmental defects

**DOI:** 10.1101/2021.04.02.438214

**Authors:** Steven Fanara, Marie Schloesser, Marc Hanikenne, Patrick Motte

**Affiliations:** InBioS-PhytoSystems, Functional Genomics and Plant Molecular Imaging and Centre for Assistance in Technology of Microscopy (CAREm), University of Liège, 4000 Liège, Belgium

## Abstract

The plant SR (serine/arginine-rich) splicing factor SR45 plays important roles in several biological processes, such as splicing, DNA methylation, innate immunity, glucose regulation and ABA signaling. A homozygous Arabidopsis *sr45-1* null mutant is viable, but exhibits diverse phenotypic alterations, including delayed root development, late flowering, shorter siliques with fewer seeds, narrower leaves and petals, and unusual numbers of floral organs. Here, we report that the *sr45-1* mutant presents an unexpected constitutive iron deficiency phenotype characterized by altered metal distribution in the plant. RNA-Sequencing highlighted severe perturbations in metal homeostasis, phenylpropanoid pathway, oxidative stress responses, and reproductive development. Ionomic quantification and histochemical staining revealed strong iron accumulation in the *sr45-1* root tissues accompanied by an iron starvation in aerial parts. We showed that some *sr45-1* developmental abnormalities can be complemented by exogenous iron supply. Our findings provide new insight into the molecular mechanisms governing the phenotypes of the *sr45-1* mutant.

**One sentence summary:** The *sr45-1* mutation affects Fe homeostasis, which results in reproductive defects

## Introduction

The precursor messenger RNA (pre-mRNA) splicing process is a crucial step in the regulation of gene expression. Splice-site selection is carried out by the binding of many *trans*-acting factors including the serine/arginine-rich (SR) splicing factors (Meyer *et al*., 2015; Jeong, 2017). Alternative splicing (AS) of a specific pre-mRNA can lead to the synthesis of multiple mRNAs and affects up to 70% of multi-exon genes in Arabidopsis (*Arabidopsis thaliana*) (Chamala *et al*., 2015). AS allows plants to cope with environmental challenges by modulating gene expression and biological processes (Palusa *et al*., 2007; Palusa and Reddy, 2010; Shang *et al*., 2017). SR proteins belong to a highly conserved family in multicellular eukaryotes and are characterized by a modular structure consisting of one or two N-terminal RNA-binding domains, called RNA recognition motifs (RRMs), and a C-terminal domain rich in arginine-serine dipeptide repeats (RS) mainly involved in protein-protein interactions (Barta *et al*., 2010; Manley and Krainer, 2010; Califice *et al*., 2012). In Arabidopsis, nineteen SR proteins constitute seven subfamilies according to their specific modular organization (Califice *et al*., 2012). Among these, the atypical SR45 protein contains two RS domains flanking a central and unique RRM (Golovkin and Reddy, 1999). The *sr45-1* null mutant exhibits some developmental defects, including delayed root development, late flowering, shorter siliques with fewer seeds, narrower leaves and petals, and unusual numbers of floral organs (Ali *et al*., 2007; Zhang *et al*., 2017). *SR45* undergoes alternative splicing producing two isoforms: the isoform *SR45.1* diverges from *SR45.2* by eight amino acids due to the presence of an additional 21-nucleotides sequence (Zhang and Mount, 2009). These isoforms fulfil distinct roles during Arabidopsis development, since *SR45.1* restores exclusively flowers defect and *SR45.2* only complements the delayed root development when expressed in the *sr45-1* background (Zhang and Mount, 2009).

SR45 was demonstrated to regulate the splicing of functionally diverse targets, thereby acting in ABA signaling and plant defense (Carvalho *et al*., 2010; Xing *et al*., 2015; Zhang *et al*., 2017). SR45 can act as a negative regulator of glucose and ABA signaling during early seedling development by modulation of SnRK1.1 stability through regulation of *5PTase13* splicing under glucose treatment (Carvalho *et al*., 2010; Carvalho *et al*., 2016). Knockout mutants for *RS40*, *RS41* and *SR45* all displayed ABA hypersensitivity (Carvalho *et al*., 2010; Chen *et al*., 2013). The *sr34b* mutation leads to the stabilization of an alternatively spliced *<u>I</u>RON-<u>R</u>EGULATED <u>T</u>RANSPORTER <u>1</u>* (*IRT1*) mRNA. Accumulation of the IRT1 divalent cation transporter in turn induces an increased uptake of cadmium (Cd) ions into the root, hence a hypersensitivity to the toxic metal Cd (Zhang *et al*., 2014). Mutations of several *SR* factors in rice (*Oryza sativa*) also depict the critical role played by alternative splicing in plant response to mineral nutrient status (Dong *et al*., 2018). Plant SR proteins actively participate to the regulation of alternative splicing under abiotic stress (Laloum *et al*., 2018; Albaqami *et al*., 2019).

Iron (Fe) is an essential micronutrient and cofactor responsible for redox state modulation (Nouet *et al*., 2011; Thomine and Vert, 2013). Since an overproduction of reactive oxygen species (ROS) may result from Fe overaccumulation (Ravet *et al*., 2009), a physiological balance of Fe uptake and accumulation has to be controlled and maintained in plant. Dicots such as Arabidopsis rely on a three-step reduction-based strategy to acquire Fe (Römheld and Marschner, 1986). The H^+^-ATPase AHA2 mediates proton extrusion, which leads to local soil acidification and Fe solubilization (Santi and Schmidt, 2009). With the help of chelators, such as coumarin phenolic compounds (scopoletin, fraxetin and sideretin) that greatly facilitate the acquisition of solubilized Fe from soils (especially at high pH ≥ 7) (Mladenka *et al*., 2010; Fourcroy *et al*., 2014; Schmidt *et al*., 2014; Rajniak *et al*., 2018; Siwinska *et al*., 2018; Tsai *et al*., 2018), the enzyme <u>F</u>ERRIC <u>R</u>EDUCTION <u>O</u>XYDASE <u>2</u> (FRO2) catalyzes the reduction of ferric [Fe(III)] to ferrous [Fe(II)] Fe (Robinson *et al*., 1999). The import of the ferrous Fe ions into root cells is finally performed by IRT1 (Vert *et al*., 2002; Castaings *et al*., 2016). Those three major components, as well as transcription factors from the myeloblastosis (MYB) family (MYB10 and MYB72) and several Fe homeostasis genes, are under the positive regulation of the basic helix-loop-helix (bHLH) transcription factor <u>F</u>ER-LIKE IRON DEFICIENCY <u>I</u>NDUCED <u>T</u>RANSCRIPTION FACTOR (FIT) (Colangelo and Guerinot, 2004; Jakoby *et al*., 2004; Ivanov *et al*., 2012; Sivitz *et al*., 2012). While <u>E</u>THYLENE <u>IN</u>SENSITIVE<u>3</u> (EIN3) and <u>E</u>THYLENE <u>I</u>NSENSITIVE3-<u>L</u>IKE<u>1</u> (EIL1) promote its stability upon Fe deficiency (Lingam *et al*., 2011), ZAT12, BTSL1, BTSL2 and MYC2 target FIT for proteasomal degradation to avoid intake of potentially toxic metals by IRT1 upon prolonged Fe deficiency (Le *et al*., 2016; Cui *et al*., 2018; Rodríguez-Celma *et al*., 2019). Together with its paralog MYB10, MYB72 regulates the expression of two <u>N</u>ICOTIANAMINE <u>S</u>YNTHASE, *NAS2* and *NAS4*, in order to facilitate the Fe redistribution in the plant by controlling the biosynthesis of the Fe chelator NA (Palmer *et al*., 2013). Another bHLH (<u>P</u>OPE<u>YE</u>, PYE) is also transcriptionally involved in Fe redistribution, especially through the expression regulation of *NAS4* and *ZIF1*, a vacuolar nicotianamine transporter (Long *et al*., 2010; Haydon *et al*., 2012). A RING E3 ubiquitin ligase, <u>B</u>RU<u>T</u>U<u>S</u> (BTS), compromises the Fe root-to-shoot translocation network controlled by PYE by targeting PYE-interacting transcriptional co-regulator for 26S proteosomal degradation (Kobayashi *et al*., 2013; Selote *et al*., 2015; Hindt *et al*., 2017). The root-to-shoot translocation of Fe involves the loading of citrate and Fe in the xylem respectively by the citrate efflux transporter <u>F</u>ERRIC <u>R</u>EDUCTASE <u>D</u>EFECTIVE 3 (FRD3) (Durrett *et al*., 2007) and possibly by Ferroportin 1 (FPN1) (Morrissey *et al*., 2009). Since endodermis suberization alters the uptake of nutrient, suberization is particularly delayed in plants growing under Fe deficiency (Baxter *et al*., 2009; Geldner, 2013; Kamiya *et al*., 2015; Barberon *et al*., 2016). This response is tightly regulated by the ethylene and ABA stress hormones which are, respectively, suppressor and activator of suberization and thus are positive and negative regulators of the root Fe-uptake, respectively (Barberon *et al*., 2016).

Alternative splicing is essential to ensure metal tolerance in plants as seen in *sr* rice mutants (Zhang *et al*., 2014; Dong *et al*., 2018). A case example is the *<u>Z</u>INC-<u>I</u>NDUCED <u>F</u>ACILITATOR <u>2</u>* (*ZIF2*) gene that undergoes intron retention in its 5’UTR to promote zinc tolerance through enhancement of its translation (Remy *et al*., 2014). Developmental alterations in the *sr45-1* mutant are strikingly similar to phenotypes associated to altered metal homeostasis, such as shorter roots and shorter siliques with fewer seeds as observed in *frd3-7* or *opt3-2* mutants (Stacey *et al*., 2008; Roschzttardtz *et al*., 2011). Although many genes involved in root responses to Fe deficiency have been identified, much less is known about the importance of their post-transcriptional processing. In this study, we explored the contribution of SR45 to Fe homeostasis in Arabidopsis roots. We show that the vegetative development of *sr45-1* plants is greatly impacted under Fe deficiency. Upon Fe supply, roots are slightly shorter because of Fe accumulation in the vascular system and the production of an oxidative burst. Fe is less concentrated in the aerial part of the mutant compared to wild-type because of impaired root-to-shoot translocation. We performed RNA-Sequencing (RNA-Seq) on wild-type and *sr45-1* roots upon Fe deficiency and control condition, which demonstrated dramatic transcriptional changes of genes involved in Fe homeostasis, ROS responses and reproductive development. As a consequence, a local metal imbalance appears in reproductive tissues of *sr45-1*, leading to shorter siliques, reduced seed number per silique, and smaller and narrower seeds. These phenotypes are fully restored in mutant plants upon exogenous Fe supply, revealing a connection between Fe uptake and developmental defects of *sr45-1*.

## Results

### The “Metal ion transport” GO category is enriched among SR45-associated RNAs

RNA-precipitation experiments recently conducted on seedlings and inflorescences revealed that SR45 binds to and regulates functionally diverse RNAs referred to as <u>S</u>R45-Associated <u>R</u>NAs (SAR) (Xing *et al*., 2015; Zhang *et al*., 2017). Here, we submitted a merged set of SAR (7799 genes) to a functional enrichment meta-analysis to identify the putative processes and pathways controlled by SR45 **(Supplemental Table S1)**: 404 Gene Ontology (GO) categories were significantly (*p*-value < 0.05) enriched among the SR45 targets and consistently illustrated the previously described phenotypes of the *sr45-1* mutant. For instance, the involvement of SR45 in salt tolerance (Albaqami *et al*., 2019) was reflected in numerous biological processes related to ion transport. Surprisingly, a biological process called “metal ion transport” (116 genes) was significantly enriched, within which several sub-GO enrichments related to divalent cation and iron transport were further observed **(Supplemental Table S1)**. Moreover, 38.5% of SAR corresponded to genes identified in transcriptomic studies as differentially expressed upon varying Fe supply (hereinafter referred as “iron-responsive genes”) **(Supplemental Figure S1)**.

### Metal levels are increased in the roots of *sr45-1*

These observations led us to examine whether the *sr45-1* mutation results in any Fe homeostasis perturbation and therefore the ionome was profiled in tissues of both *sr45-1* and WT (Col-0) plants grown in Fe-deficient (0 µM) and Fe-sufficient (10 µM) conditions. In Fe-sufficient condition, the roots of *sr45-1* accumulated more Fe (136%), Mn (218%) and Zn (150%) compared to the WT. In contrast, Cu levels were similar in the two genotypes **(Figure 1A-D)**. Upon Fe deficiency, Fe and Zn accumulated similarly in *sr45-1* and WT roots, but *sr45-1* roots accumulated more Cu (125%) and Mn (146%). Conversely, *sr45-1* accumulated less Fe (52%), Mn (38%), Zn (37%) and Cu (33%) than the WT in shoots under Fe deficiency **(Supplemental Figure S2A-D)**. Accumulation reduction of Fe (17%), Mn (18%) and Zn (32%) was also significant in *sr45-1* shoots under Fe-sufficient condition **(Supplemental Figure S2A and C-D)**. While root macronutrients (Ca, K and Mg) levels were similar between genotypes under Fe deficiency and sufficiency, Ca levels were reduced by 26% in mutant shoots at 0 µM Fe but K levels were increased by 36% at 10 µM Fe, and no significant changes were observed in mutant shoots regarding Mg levels **(Supplemental Figure S3A-C)**.

**Figure 1.**
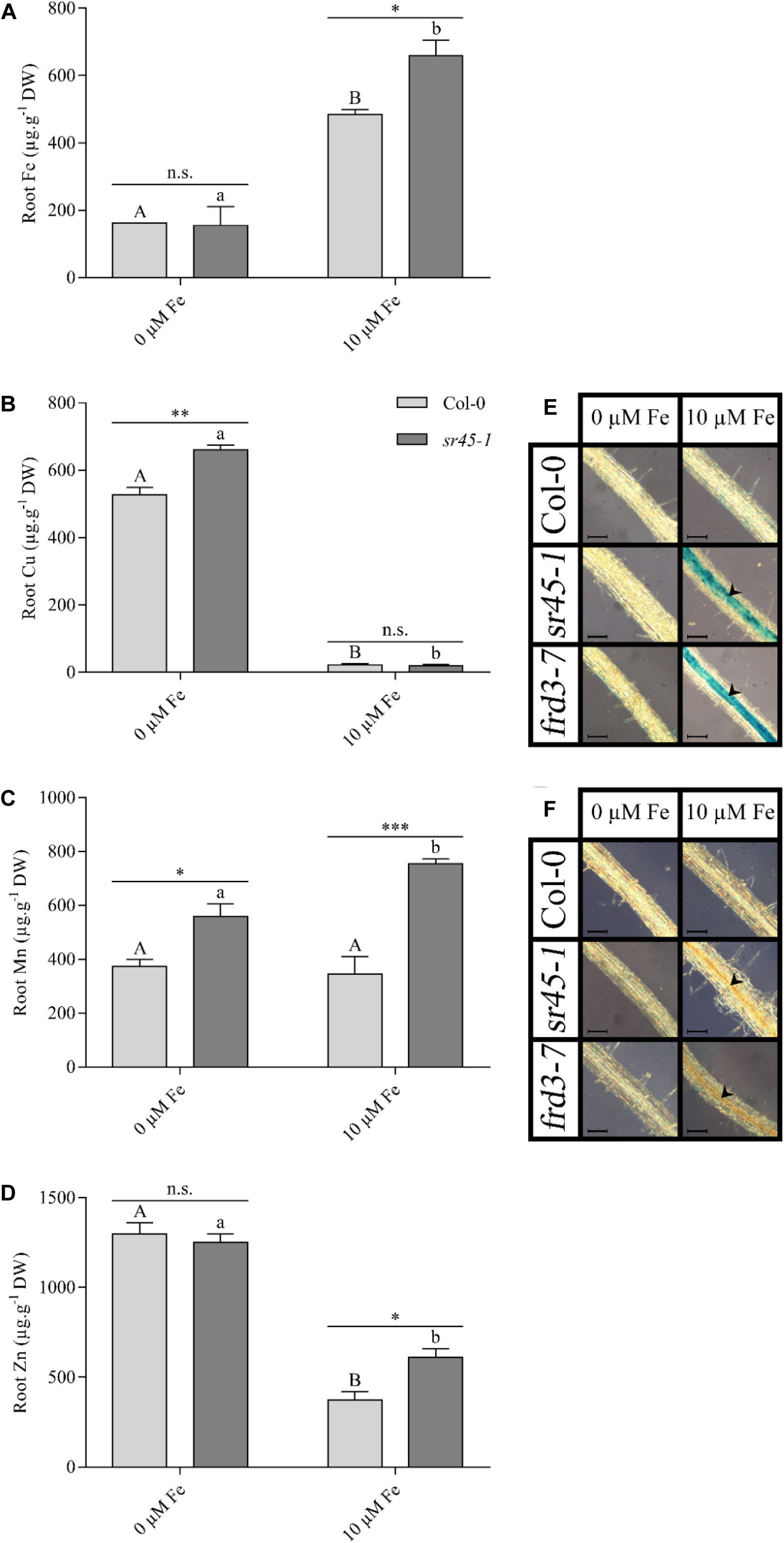
Metal concentration, iron staining and H_2_O_2_ staining in roots upon iron deficiency and iron excess. **(A)** Iron (Fe), **(B)** copper (Cu), **(C)** manganese (Mn) and **(D)** zinc (Zn) concentrations in roots of wild-type (Col-0) and mutant (*sr45-1*) plants grown hydroponically in Hoagland medium supplemented with 0 (deficiency) or 10 (control) µM Fe. Values represent means ± SEM (from one experiment representative of two independent experiments, each including 2 or 3 series of 2 plants per genotype and condition). Data were analyzed by two-way ANOVA followed by Bonferroni multiple comparison post-test. Statistically significant differences between means between genotypes are indicated by stars (* P<0.05, ** P<0,01, *** P<0.001) or between treatments within genotypes by different letters (P<0.05). n.s.: not significant. **(E)** Iron accumulation visualized using Perls staining in roots of Col-0 and *sr45-1*. **(F)** H_2_O_2_ accumulation stained using DAB in roots of Col-0 and *sr45-1*. The pictures are representative of 2 independent experiments. Arrowheads show Fe or H_2_O_2_ accumulation in roots. Scale bar: 100 μm.

In Fe sufficiency condition, Perls’ staining revealed strong Fe accumulation in the vascular cylinder of *sr45-1* roots, **(Figure 1E)**, a phenotype strikingly similar to a *ferric reductase defective3* (*frd3*) loss-of-function mutant (Green and Rogers, 2004; Roschzttardtz *et al*., 2009; Scheepers *et al*., 2020). Because of this phenotypic similarity, we decided to concomitantly analyse *frd3-7* as a control for defective Fe homeostasis **(Figure 1E)**. No Fe accumulation in root vascular tissues occurred upon Fe deficiency in the three genotypes. It is well described that overaccumulation of Fe can trigger ROS overproduction, ultimately leading to oxidative stress (Reyt *et al*., 2015). Fe and ROS accumulation were shown to correlate in root vascular tissues of *frd3-7* (Scheepers *et al*., 2020). DAB staining, revealing H_2_O_2_ accumulation, was accordingly detected in the central vasculature of the root in both *sr45-1* and *frd3-7* mutants but not in the WT **(Figure 1F)**. Altogether, the Fe accumulation pattern in roots and shoots of *sr45-1* plants are indicative of a defective metal root-to-shoot translocation of Fe in *sr45-1*.

### The *sr45-1* mutant is sensitive to the Fe status

Based on these observations, we investigated the effect of Fe supply on the *sr45-1* mutant development by growing either seedlings for 14 days on Fe-depleted (0 µM Fe), control (10 µM Fe) or supplemented with Fe excess (50 µM Fe) agar solid Hoagland or 6-week-old plants in hydroponic solution in Fe-deficient and Fe-sufficient conditions **(Figure 2**, **Supplemental Figure S4A)**. In seedlings, the root growth of the *sr45-1* mutant was reduced by 47% under Fe deficiency and 68% under Fe excess compared to WT **(Supplemental Figure S4A)**. The reduction of root biomass correlated with a reduction of shoot biomass with a loss of 46%, 38.5% and 21% at Fe deficiency, sufficiency and excess, respectively, compared to WT at Fe sufficiency **(Supplemental Figure S4B)**.

**Figure 2.**
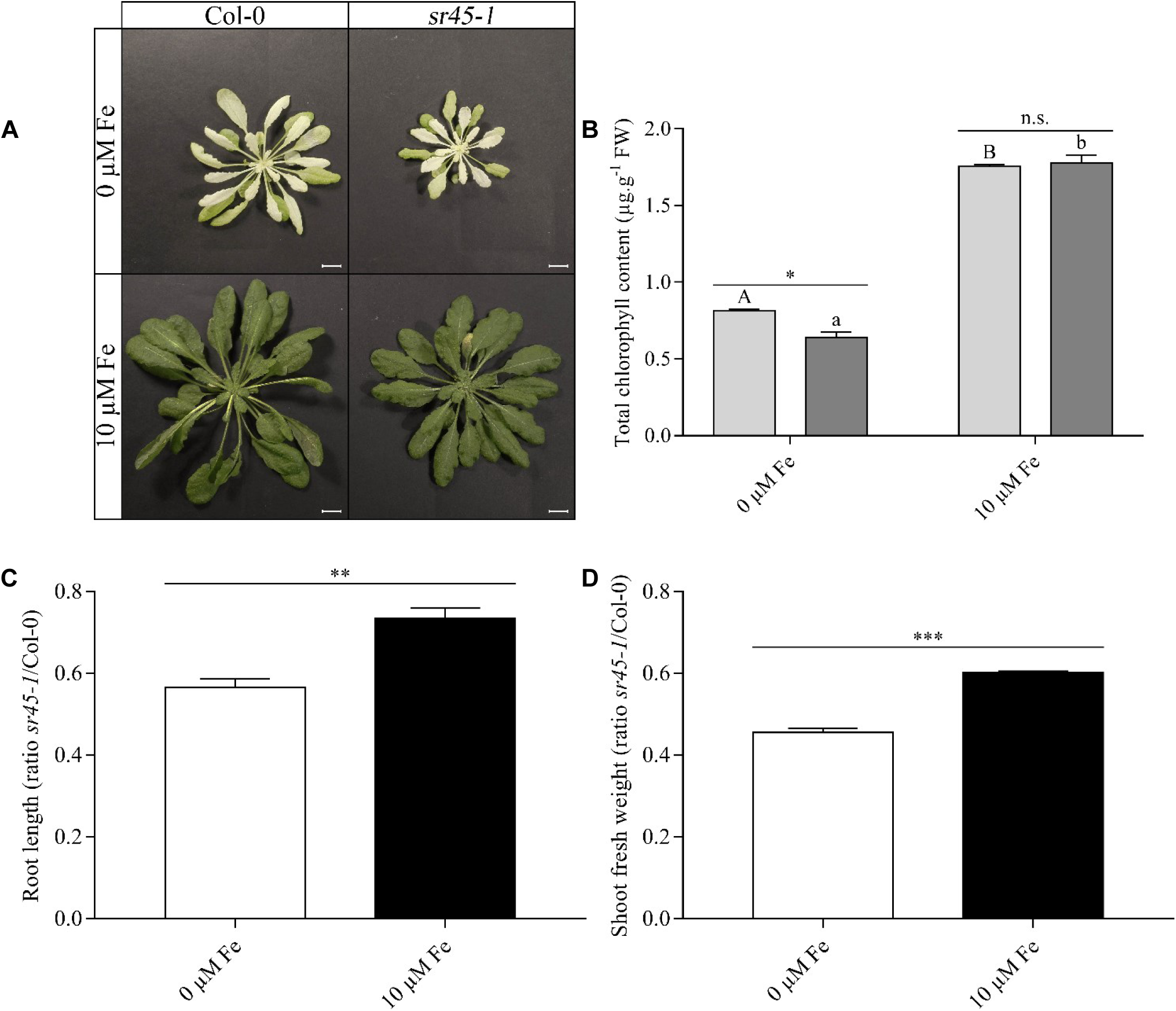
Phenotypes of wild-type and *sr45-1* plants upon iron deficiency. **(A)** Representative pictures and **(B)** quantification of total chlorophyll content of wild-type (Col-0) and mutant (*sr45-1*) plants grown hydroponically in Hoagland medium supplemented with 0 (deficiency) or 10 (control) µM Fe. Scale bar: 1 cm. **(B)** Ratios of root length and **(C)** shoot fresh weight in *sr45-1 versus* Col-0 plants. Data were analyzed by two-way ANOVA followed by Bonferroni multiple comparison post-test **(B)** or by a t-test **(C-D)**. Statistically significant differences between means between genotypes are indicated by stars (* P<0.05, ** P<0,01, *** P<0.001) or between treatments within genotypes by different letters (P<0.05). n.s.: not significant. Scale bar: 1 cm.

In adult plants, *sr45-1* root growth was significantly decreased in both deficiency and excess conditions (**Supplemental Figure S4C**). In response to Fe deficiency, roots of the WT and *sr45-1* were 27.5% and 43% shorter, respectively, compared to Fe sufficiency. Similarly, the shoot fresh weight of both wild-type and *sr45-1* mutant plants was affected by Fe deficiency with a significant biomass loss of 61.5% and 71.3%, respectively, compared to the Fe-sufficient condition **(Supplemental Figure S4D)**. Although *sr45-1* development was already delayed in control condition, the *sr45-1*/WT ratio for both root length **(Figure 2C)** and shoot fresh weight **(Figure 2D)** indicated that adult *sr45-1* plants were more sensitive to Fe deficiency than the WT. Fe deficiency resulted in chlorosis in shoots of both WT and mutant. However, the aerial parts of the *sr45-1* mutant presented a more severe chlorosis **(Figure 2A-B)** at 0 µM Fe than the WT, confirming higher sensitivity of the mutant to Fe deficiency. Exposure to higher Fe supply (50 µM) additionally suggested that the mutant had a perturbed root-to-shoot Fe translocation: whereas the WT suffered of Fe toxicity with less biomass production in aerial parts in these conditions (significant loss of 34%), higher Fe supply did not cause further delay nor improved shoot growth of adult *sr45-1* plants **(Supplemental Figure S4D)**. This confirmed the impaired Fe root-to-shoot distribution and inferred that increasing Fe supply would trigger toxicity in *sr45-1* roots.

### Zinc accumulation in roots of *sr45-1* is toxic

The 2-fold and 1.5-fold increases in accumulation of respectively Mn and Zn in *sr45-1* roots upon Fe sufficiency **(Figure 1C-D)** may cause growth reduction as previously observed (Dučić and Polle, 2007; Lei *et al*., 2007; Kawachi *et al*., 2009; Fukao *et al*., 2011; Shanmugam *et al*., 2011; Marschner and Marschner, 2012; Millaleo *et al*., 2013; Scheepers *et al*., 2020). Thus, we investigated the influence of those metals on the growth of *sr45-1*. Contrary to Mn depletion, Zn depletion did not significantly impact the root growth of the WT in our growth condition on agar plates. In contrast, root length of *sr45-1* was significantly longer by 12% upon Zn deficiency, but roots remained 50% smaller compared to WT in control condition (**Figure 3**), suggesting that Zn accumulation in roots was toxic and indeed partially contributed to the growth defect of *sr45-1*. Mn and Zn deficiency did not influence Fe or H_2_O_2_ accumulation in the *sr45-1* root vasculature **(Supplemental Figure S5A-B)**.

**Figure 3.**
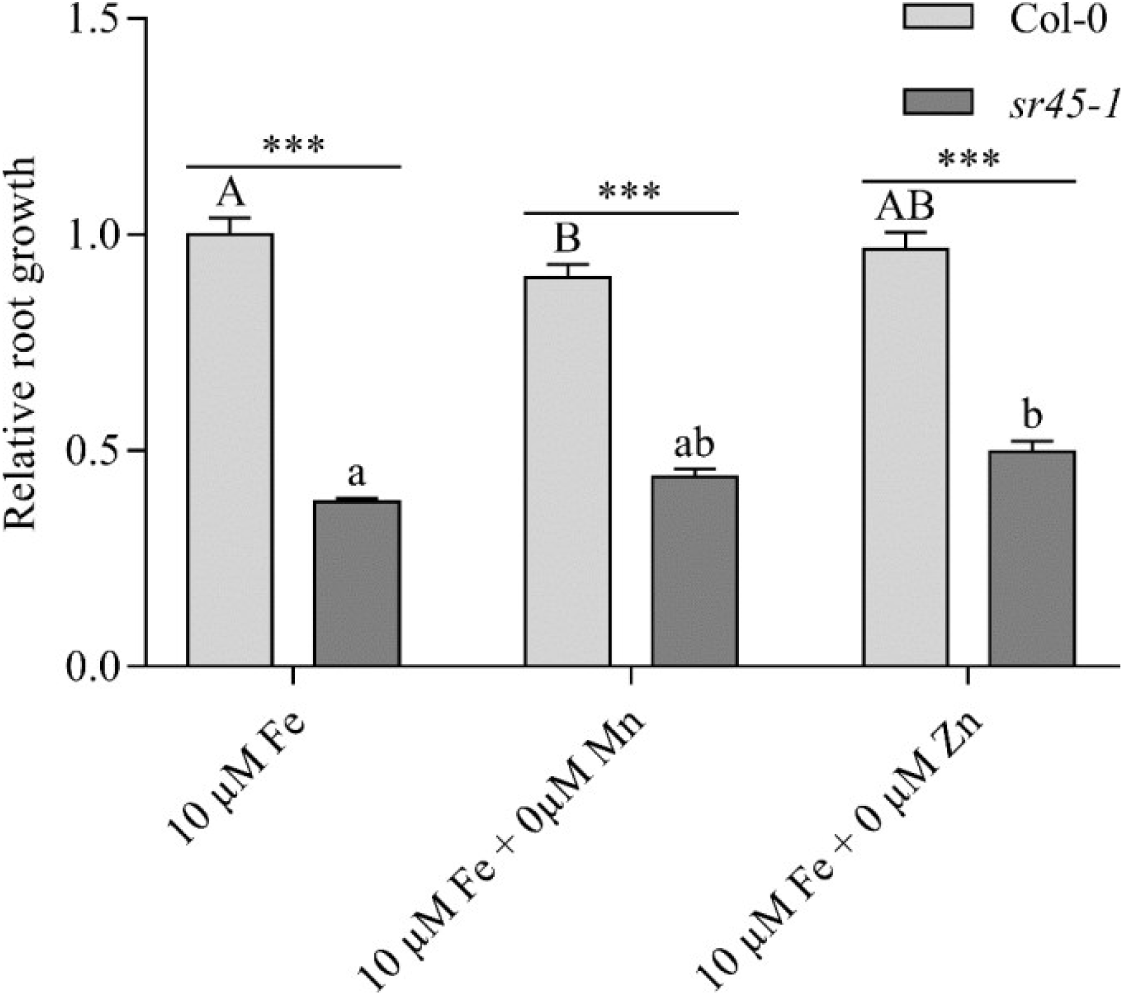
Toxicity of manganese and zinc in *sr45-1* roots. Relative root length of wild-type (Col-0) and mutant (*sr45-1*) plants grown vertically in petri dishes containing Hoagland medium supplemented with iron (10 µM Fe-HBED) but not manganese (0 µM MnSO_4_) or zinc (0 µM ZnSO_4_). Root growth is relative to Col-0 at 10 µM Fe-HBED, 5 µM MnSO_4_ and 1 μM ZnSO_4_. Values represent means ± SEM (from four independent experiments, each including 2 series of 6 seedlings per genotype and condition). Data were analyzed by two-way ANOVA followed by Bonferroni multiple comparison post-test. Statistically significant differences between means between genotypes are indicated by stars (*** P<0.001) or between treatments within genotypes by different letters (P<0.05).

The shoot fresh weight of *sr45-1*, which was lower by 28% in control condition compared to the WT, was increasingly affected by Mn and Zn deficiencies, leading to a biomass loss of 70% and 73.5%, respectively, compared to the WT in Fe sufficiency **(Supplemental Figure S5C)**. In Zn-deficient condition, Col-0 shoot biomass was decreased by 68.5%, resulting in the absence of significant changes between both genotypes. Altogether, these results pointed to a Zn toxicity in roots and to a defective metal homeostasis, possibly in acquisition and mobilization of metals.

### Transcriptome profiling of *sr45-1* mutant roots

To uncover the molecular mechanisms underlying metal distribution in *sr45-1*, a transcriptomic profiling of WT and *sr45-1* root tissues of adult plants grown hydroponically in Fe-deficient (0 µM) and Fe-sufficient (10 µM) conditions was conducted using RNA-Seq. A limited number of differentially expressed genes (DEGs) [fold change ≥ 2 or ≤ −2 and false discovery rate (FDR) < 0.05] were identified between WT and mutant roots in Fe-sufficient (29 DEGs) and Fe-deficient conditions (49 DEGs) **(Figure 4 and Supplemental Table S2)** and no enriched GO functional category was identified among either of those sets of DEGs **(Supplemental Table S3)**. *sr45-1* mutant plants presented less transcriptomic changes in response to Fe deficiency than Col-0 (1350 *vs* 2442 DEGs) and accordingly, the number of enriched GO biological processes was considerably reduced in the mutant (33 *vs* 50 and 20 *vs* 75 for the up- and down-regulated genes, respectively) **(Supplemental Table S3)**. In fact, in response to Fe deficiency, Col-0 displayed extensive changes in the expression of genes contributing to diverse biological processes that were not over-represented in the mutant roots. For instance, the response of Col-0 roots to Fe deficiency involved the induction of metal ion homeostasis and transport, as well as the repression of several functions including phenylpropanoid biosynthetic process, cell wall modifications, lignin biosynthesis, biotic and hormonal responses **(Supplemental Table S3)**. The molecular alteration in the *sr45-1* mutant roots under Fe deficiency included perturbation in cellular iron ion homeostasis (*FER* and *VTL* genes) and oxidative stress response (through down-regulation of, respectively, 7 and 35 genes), as well as in anion transport and sulfate transport (through up-regulation of, respectively, 34 and 6 genes). Surprisingly, while response to ethylene was a significant over-represented function in both Col-0 and *sr45-1* roots upon Fe deficiency, divergence appeared: 42 genes were under-expressed in Col-0 (against 9 of them in *sr45-1*) and 22 genes were over-expressed in the mutant (against 18 of them in Col-0) **(Supplemental Table S3)**.

**Figure 4.**
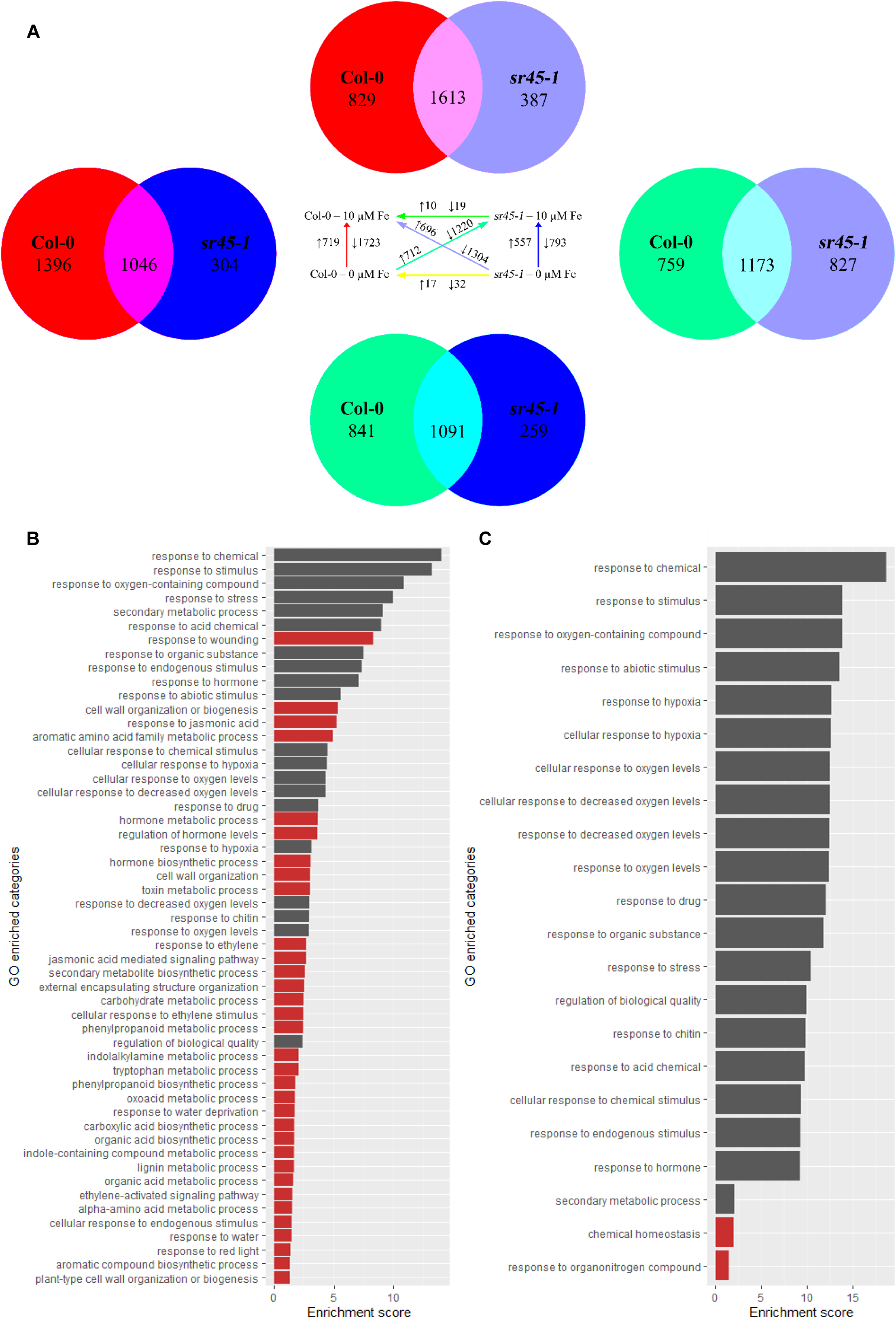
Transcriptomic analysis in roots upon iron deficiency and sufficiency. **(A)** Summary statistics of up- (↑) or down-regulated (↓) genes in different pair-wise comparisons. The direction of comparisons is indicated by arrows. Overlaps in Venn diagrams represent deregulated genes in both Col-0 and *sr45-1* roots in respective comparisons (indicated by colors). Right circle: Number of differentially regulated genes solely associated to *sr45-1* roots (SDRs). Left circle: Number of differentially regulated genes solely associated to Col-0 (CDRs). Bar graphs of Gene Ontology (GO) enriched categories in differentially expressed CDR genes **(B)** and SDR genes **(C)**. The enrichment score represents –log_10_(*p*-value). Dissimilar GO terms between genotypes are colored in red (see also **Supplemental Table S5**).

To further characterize *sr45-1* root transcriptome and fine-tune our understanding of the metal homeostasis regulated by SR45, additional comparisons were performed to discover SR45 differentially regulated genes (SDR; DEGs significantly and specifically deregulated in *sr45-1* roots in respective comparison) **(Figure 4)**: SDR included 304 and 387 in the comparison of the Fe deficiency DEGs in the WT (2442 genes) with either (i) the Fe deficiency DEGs in the mutant (1350 genes) or (ii) the DEGs between mutant at Fe deficiency and the WT in control condition (2000 genes), respectively. The reciprocal of the latest comparison yielded 259 additional SDR. Finally, 827 SDR were obtained when intersecting the DEGs obtained when comparing a genotype (WT or *sr45-1*) in deficiency with the other genotype in sufficiency. Together with the DEGs identified between genotypes in Fe-sufficient (29) and Fe-deficient (49) conditions, it represented 1071 unique SDR compared to 1844 CDR (Col-0 differentially regulated genes) **(Supplemental Table S4)**. The analysis of GO enrichment highlighted molecular contribution of both CDRs and SDRs to many similar biological pathways. However, CDRs included genes involved in biological functions that were not over-represented in the mutant, such as response to jasmonic acid and ethylene, cell wall organization, phenylpropanoid biosynthetic and metabolic processes or lignin metabolic process **(Figure 4B**, **Supplemental Table S5)**. SDRs contribute additionally to two biological functions unrepresented in CDRs, namely chemical homeostasis and response to organonitrogen compound, which include genes involved in Fe, Mn, Zn, Cu and Cd homeostasis **(Figure 4C**, **Supplemental Table S5)**.

These observations gave evidence that the mutant is pervasively affected in many pathways upon Fe deficiency. It raised the question whether *sr45-1* mutant plants were able to accurately regulate the response of key genes or whether adult mutant plants were coping with residual induction of the metal homeostasis due to defective seedlings (as shown by the severe toxic symptoms displayed in response to Fe supply **(Supplemental Figure S4)**.

### The Fe acquisition machinery is constitutively induced in *sr45-1* seedlings

As the ionome and the growth of the mutant were affected by the Fe status (**Figure 1** and **Supplemental Figure S4**), gene expression (using qRT-PCR) and protein activity of key components of the Fe uptake machinery were examined in roots of 14-day-old seedlings, as well as 9-week-old roots grown in Fe deficiency and sufficiency conditions. The expression of the bHLH transcription factor *FIT* was up-regulated by Fe deficiency in both WT and *sr45-1* seedling roots compared to the 10 µM Fe condition. However, *FIT* expression was ∼47-51% higher in *sr45-1* seedlings compared to Col-0 in both Fe-sufficient and Fe-deficient conditions **(Figure 5A)**. Similarly, the FIT targets *AHA2*, *FRO2* and *IRT1* were also more highly expressed (124-208%) in *sr45-1* roots in both conditions compared to Col-0 **(Figure 5B-D)**, consistent with increased acidification of the medium (mostly driven by AHA2) **(Supplemental Figure S6A)** and ferric chelate reductase activity (mostly driven by FRO2) in *sr45-1* roots **(Supplemental Figure S6B)**. Altogether, these observations were indicative of a constitutive Fe deficiency response in *sr45-1* roots upon control Fe supply (10 µM). This response was further aggravated when no Fe (0 µM) was supplied.

**Figure 5.**
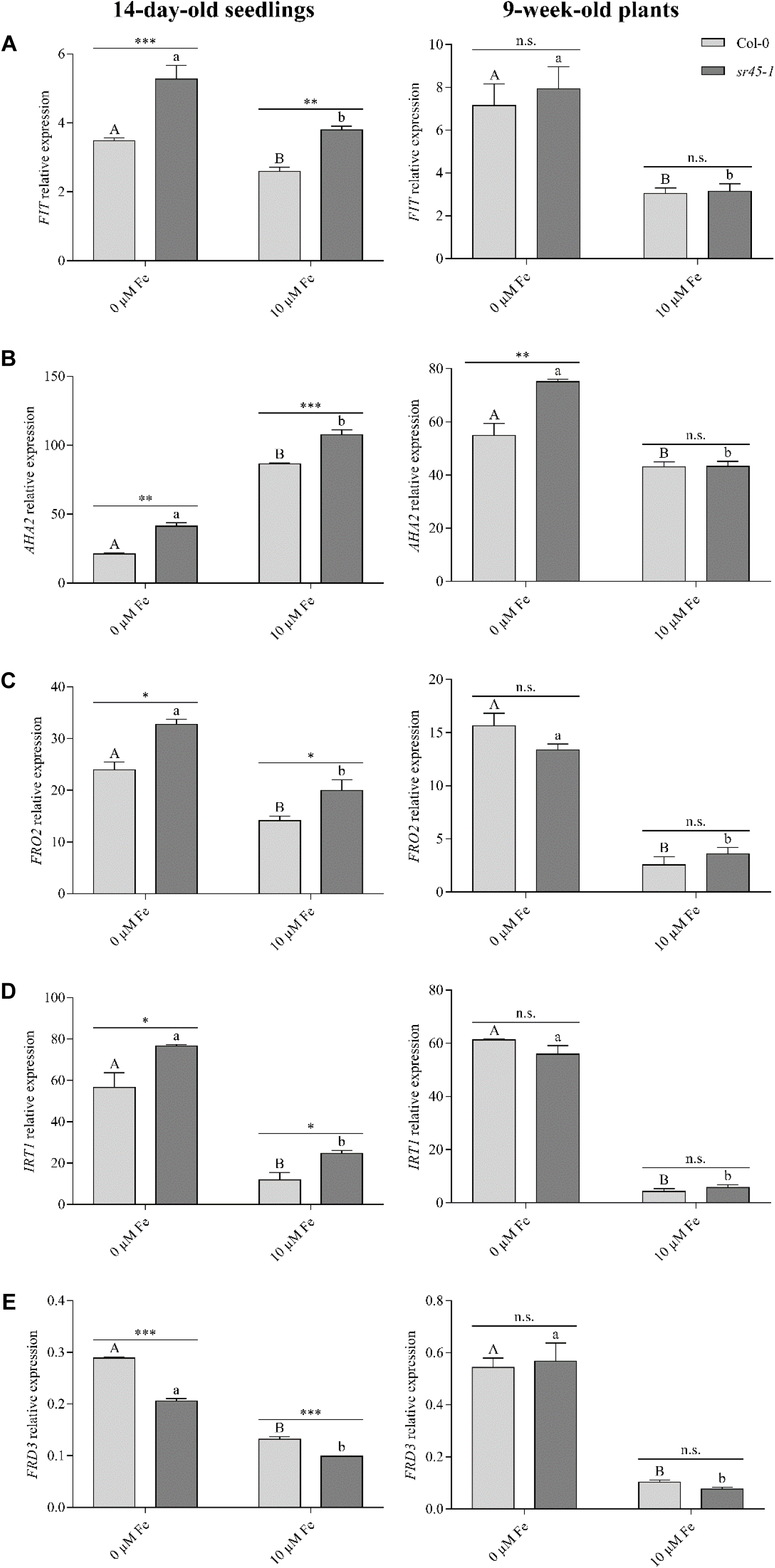
Fe uptake machinery in *sr45-1* seedlings and adult plants. Quantitative RT-PCR analysis of expression of **(A)** *FIT*, **(B)** *AHA2*, **(C)** *FRO2*, **(D)** *IRT1* and **(E)** *FRD3* genes wild-type (Col-0) and mutant (*sr45-1*) seedlings grown vertically in petri dishes containing Hoagland medium supplemented with 0 or 10 μM iron (Fe) **(left panel)** and in roots of wild-type (Col-0) and mutant (*sr45-1*) plants grown hydroponically in Hoagland medium supplemented with 0 or 10 µM Fe (RNA-seq samples) **(right panel)**. Values represent means ± SEM (from four biological replicates, each consisting of 2-6 plants per genotype and condition) and are relative to *At1g58050*. Data were analyzed by two-way ANOVA followed by Bonferroni multiple comparison post-test. Statistically significant differences between means between genotypes are indicated by stars (* P<0.05, ** P<0,01, *** P<0.001) or between treatments within genotypes by different letters (P<0.05). n.s.: not significant.

In adult roots, all key components of the Fe uptake machinery were induced in both Col-0 and *sr45-1* upon Fe deficiency, but no significant deviation was observed between genotypes for *FIT*, *FRO2* or *IRT1* expression. However, *AHA2* expression was ∼36.8% higher in *sr45-1* roots compared to Col-0. While *FIT* expression was not induced in adult mutant roots at Fe deficiency or sufficiency **(Figure 5A)**, the expression of genes encoding the MYB72, bHLH38, bHLH39, bHLH100 and bHLH101 transcription factors, which are known to act in conjunction with FIT to induce the Fe acquisition machinery (*AHA2*, *FRO2* and *IRT1*) (Yuan *et al*., 2008; Palmer *et al*., 2013; Wang *et al*., 2013), was significantly up-regulated in the *sr45-1* mutant under Fe deficiency compared to Col-0 **(Supplemental Figure S7)**.

Since Fe accumulation in the root vasculature was observed in both *sr45-1* and *frd3-7* mutants, the relative expression level of *FRD3* in *sr45-1* roots was also analyzed. The level of *FRD3* was reduced by 29% and 25% in the mutant upon Fe-deficient and Fe-sufficient conditions, respectively **(Figure 5E)**. In agreement with this observation, both *sr45-1* and *frd3-7* accumulated a higher level of citrate, the substrate of FRD3, in roots, but it was significantly lower in *sr45-1* **(Supplemental Figure S6C)**. In contrast, adult mutant roots did not show any deviation in *FRD3* expression compared to the WT **(Figure 5E)**.

All results suggested that, contrary to adult plants, *sr45-1* seedlings display a constitutive iron deficiency response independently of the iron concentration used (0 µM or 10 µM Fe) and that the root-to-shoot Fe translocation may be impaired through down-regulation of *FRD3* in young roots.

### Influence of coumarins on the developmental defects of *sr45-1* mutant seedlings

The phenylpropanoid pathway, which participates in the synthesis of coumarins involved in Fe solubilization from the soil, including through a transcriptional control by BEE1 and MYB15 (Petridis *et al*., 2016; Chezem *et al*., 2017), both identified as SDR in this study, is improperly regulated in *sr45-1* adult roots **(Supplemental Table S3** and **Supplemental Table S5)**. In fact, many genes were down-regulated in Col-0 in response to Fe deficiency, but only few of them were affected in this fashion in the mutant **(Supplemental Figure S8)**. This suggested that Fe mobilization from the soil may be affected in the mutant. To investigate this hypothesis, Col-0 and mutant seedlings were grown on solid Hoagland media containing or not Fe-mobilizing compounds and supplemented with 10 µM Fe. A Hoagland control medium, containing Fe-HBED at pH 5.7, was used as the maximum of Fe bioavailability, and a Hoagland unavailable Fe medium, containing FeCl_3_ at neutral pH, was used as the minimum of Fe bioavailability (Tsai *et al*., 2018). Whereas the root growth of the WT was significantly reduced (∼10%) on FeCl_3_ medium, it was increased by ∼29% in the mutant **(Figure 6A)**, which confirms that Fe in a soluble form causes toxicity in *sr45-1* roots. No signal of iron accumulation nor oxidative stress was visualized in seedlings grown on the unavailable Fe medium **(Figure 6B-C)**. This suggested that *sr45-1* is less capable of mobilizing Fe from the Fe unavailable medium. We then tested whether supplementation of coumarins could help the mutant in this regard. Exogenous application of fraxetin did not affect the root growth of Col-0 nor *sr45-1* **(Figure 6A)** but led to an increase in the Fe and H_2_O_2_ accumulation in the mutant roots compared to the FeCl_3_ only condition **(Figure 6B-C)**. In the absence of Fe (0 µM FeCl_3_), no significant change in the length of WT roots was observed, but the mutant roots were significantly longer by 8.5% compared to the Fe unavailable condition, further indicating the Fe toxicity in the mutant. The further addition of fraxetin led to a significant decrease of both Col-0 (∼13.5%) and *sr45-1* (∼12.5%) root length **(Figure 6A),** whereas no Fe nor H_2_O_2_ staining was observed in seedlings **(Figure 6B-C)**. This suggests that coumarins tend to mobilize Fe (and possibly other metals) from the medium, which restores metal toxicity in the mutant.

**Figure 6.**
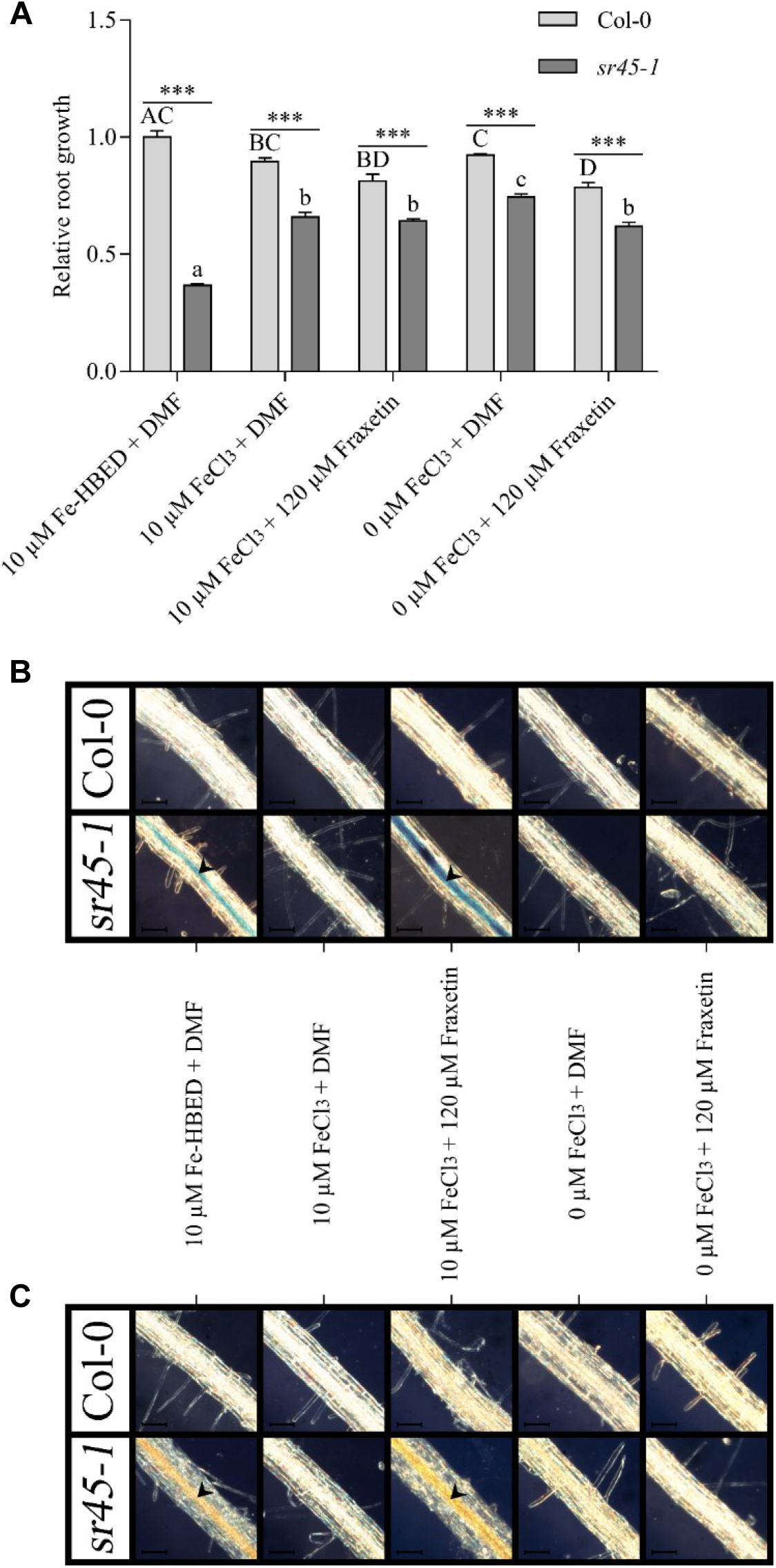
Effect of exogenous application of coumarins on *sr45-1* roots. **(A)** Relative root growth of wild-type (Col-0) and mutant (*sr45-1*) plants grown vertically in petri dishes containing Hoagland medium supplemented with 10 µM iron (Fe) at various pH (5.7 in presence of Fe-HBED or 7 in presence of FeCl_3_). Root growth is relative to Col-0 at 10 µM Fe-HBED + DMF (pH 5.7). Values represent means ± SEM (from two independent experiments, each including 3 or 4 series of 6 plants per genotype and condition). Data were analyzed by two-way ANOVA followed by Bonferroni multiple comparison post-test. Statistically significant differences between means between genotypes are indicated by stars (*** P<0.001) or between treatments within genotypes by different letters (P<0.05). **(B)** Iron accumulation and **(C)** H_2_O_2_ accumulation in roots of wild-type (Col-0) and mutant (*sr45-1*) plants grown vertically in petri dishes containing Hoagland medium supplemented with 0 (deficiency) or 10 (control) µM iron (Fe) at various pH (5.7 in presence of Fe-HBED or 7 in presence of FeCl_3_). The pictures are representative of two independent experiments. Arrowheads show Fe or H_2_O_2_ accumulation in roots. Scale bar: 100 μm.

In presence of unavailable Fe, the shoot fresh weight of Col-0 or *sr45-1* were significantly reduced by ∼33% and ∼5.5% compared to themselves in control condition **(Supplemental Figure S9A-B)**, confirming that the mutant shoots initially suffered of an iron deficiency that was not significantly aggravated in presence of unavailable Fe. Supplementation of coumarins affected the shoot fresh weight of the seedlings in the same fashion than the unavailability of Fe **(Supplemental Figure S9A-B)**, which suggests that despite the improved solubilization and uptake of Fe, the mutant did not efficiently translocate Fe from roots to shoots to ensure proper development. The total chlorophyll content of the WT or the mutant was not significantly affected in any of the tested conditions **(Supplemental Figure S9C)**.

Although the phenylpropanoid pathway is deregulated, the function of fraxetin is not defective in the mutant. In fact, the mutant is capable of remobilizing iron from the insoluble FeCl_3_ salt in presence of this coumarin, leading to root iron accumulation but not toxicity.

### Iron supplementation fully restored all the reproductive phenotypes of *sr45-1* mutant plants

We last questioned if the *sr45-1* phenotypes of shorter siliques and reduction in seed yield (Zhang *et al*., 2017) could be rescued by Fe supplementation, as shown for the *frd3-7* mutant (Roschzttardtz *et al*., 2011). Col-0 and *sr45-1* plants were therefore grown on soil until seed setting. They were daily irrigated with water without (–Fe) or with an additional treatment consisting of a weekly irrigation with sequestrene (+Fe). The sequestrene irrigation of the soil fully restored silique length **(Figure 7A-B)** and seed number per silique in the *sr45-1* mutant **(Figure 7D)**. The “shrunken” phenotype of the *sr45-1* seeds compared to Col-0, consisting of a decreased seed length, width and mass, was also fully restored upon Fe irrigation **(Figure 7C and 7E-F)**. However, Fe-irrigated mutants still displayed impaired growth of stem **(Supplemental Figure S10A)** and late flowering upon Fe treatment **(Supplemental Figure S10B-C)** (Ali *et al*., 2007). This suggested that the complementation by Fe was specific to seed development.

**Figure 7.**
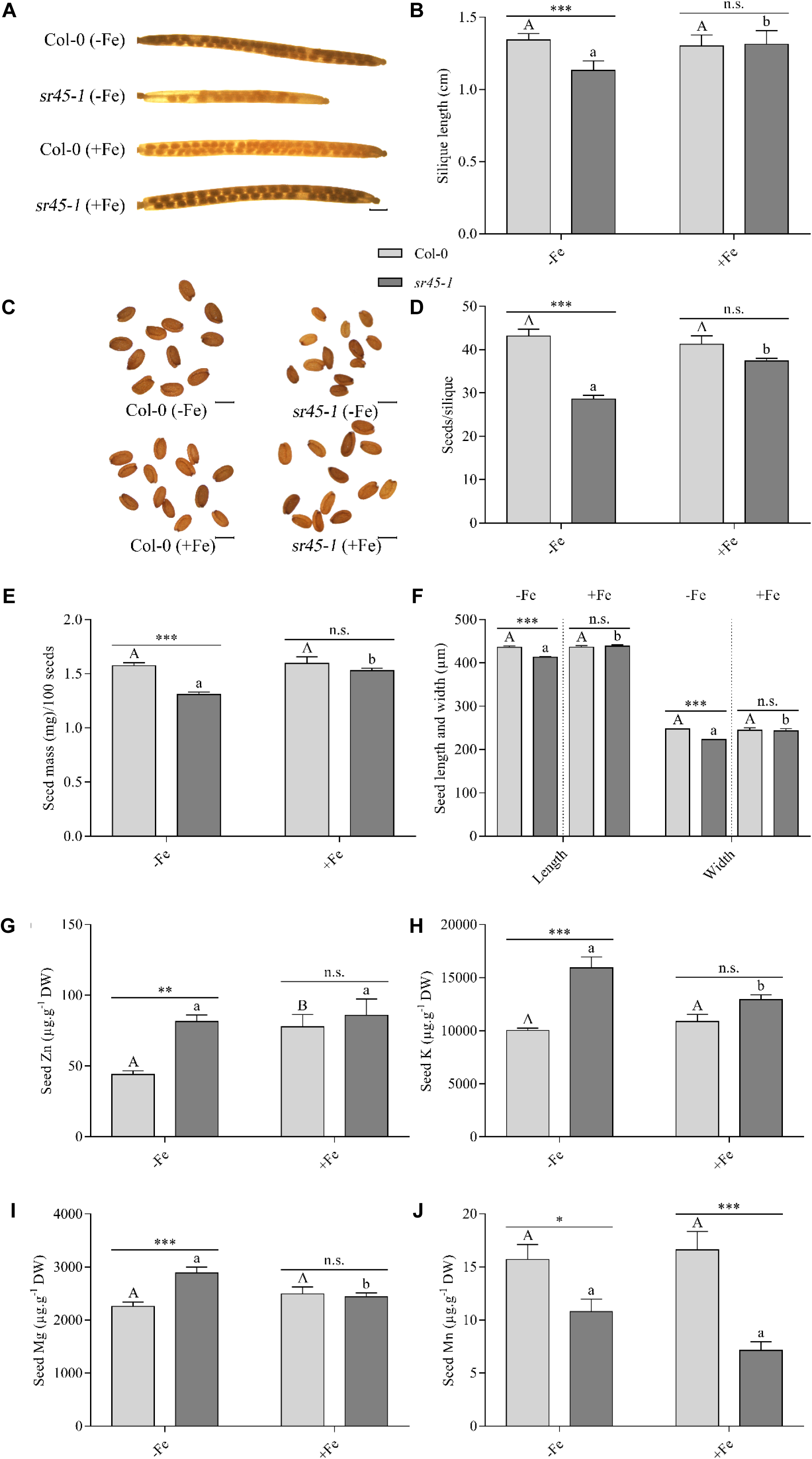
Iron irrigation fully rescues the development of the reproductive tissues. **(A)** Representative pictures of wild-type (Col-0) and mutant (*sr45-1*) siliques, **(B)** silique length, **(C)** representative pictures of wild-type (Col-0) and mutant (*sr45-1*) seeds, **(D)** number of seeds per silique, **(E)** seed mass (for pools of 100 seeds), and **(F)** seed length and width of plants grown in soil and daily irrigated with water without (–Fe) or with an additional treatment consisting of a weekly irrigation with sequestrene (+Fe). Values represent means ± SEM (from three independent experiments with 10-18 siliques pertaining to the main stem [4 plants per genotype and condition] **(B)** or with seeds obtained from 4 plants per genotype and condition **(D-F)**). Scale bars: 1 mm for siliques; 500 µm for seeds. **(G)** Zinc (Zn), **(H)** potassium (K), **(I)** magnesium (Mg) and **(HJ)** manganese (Mn) concentration in seeds (–Fe or +Fe) of wild-type (Col-0) and mutant (*sr45-1*) plants. Values represent means ± SEM (from three independent experiments with seeds obtained from 4 plants per genotype and condition). For each experiment, data were analyzed by two-way ANOVA followed by Bonferroni multiple comparison post-test. Statistically significant differences between means are indicated by stars (* P<0.05, ** P<0,01, *** P<0.001) or different letters (P<0.05). n.s.: not significant. **(F)** was obtained by performing statistical analysis for both seed length and seed width independently and then by displaying the results in a single graph.

To examine whether defects in seed morphology were linked to perturbed metal concentrations, an ionomic analysis was conducted on –Fe and +Fe seeds. Compared to WT seeds, *sr45-1* seeds displayed significantly increased in Zn, K and Mg levels but contained reduced amount of Mn. Upon Fe irrigation, these defects were completely restored for all metals but Mn **(Figure 7G-J)**. In the case of Zn, the Fe supplementation did not modulate the Zn concentration in mutant seeds but increased the Zn concentration in the one of Col-0 **(Figure 7G)**. Finally, no defect was detected in mutant seeds for micronutrients Fe and Cu, nor macronutrients Ca and P **(Supplemental Figure S10D-G)**, but Fe irrigation increased the Cu content in both WT and *sr45-1* seeds **(Supplemental Figure S10E)**. Altogether, these results suggested that the mutant seeds contained abnormal concentrations of metals and that Fe supplementation was able to restore all the ion defects (but Mn) in the mutant seeds.

## Discussion

SR45, an atypical SR protein containing two RS domains, is an auxiliary component of the spliceosome whose biological functions remained elusive (Califice *et al*., 2012). It has been recently shown that this splicing factor regulates diverse biological processes including abscisic acid signaling and innate immunity, thanks to the identification of its direct targets (SARs) in two complementary studies (Xing *et al*., 2015; Zhang *et al*., 2017). Here, combining these two datasets, identifying SARs (7799) (Xing *et al*., 2015; Zhang *et al*., 2017) and SDR genes (358) in reproductive tissues (Zhang *et al*., 2017), with an RNA-Seq analysis of the roots of the *sr45-1* mutant upon changes in Fe supply, identifying 1071 SDR genes, allowed to better describe major defects linked to the *sr45-1* mutation and hence to better depict a global function of SR45. Eighteen genes emerged as shared among SARs and SDR genes, representing probable SR45 RNA targets that are constantly deregulated regardless of the tissue and stage of development of the *sr45-1* mutant (e.g. *MYB15*, *PWD/GWD2*, *OCT3* or *AtHMP37*) **(Supplemental Table S6)**. Loss-of-function mutants of several SDR genes identified in this study display abnormal silique length and seed set, as well as seed abortion **(Supplemental Table S5 and Supplemental Figure S11)**, which are also strong phenotypes of the *sr45-1* mutant (Zhang *et al*., 2017). Among these knock-out lines, the mutation of *PWD/GWD2* (*AT4G24450*) leads to a reduced number of siliques and a decreased seed production, and *pwd/gwd2* seeds are shrunken with an irregularly shaped coat (Pirone *et al*., 2017). The *PWD/GWD2* gene is not only a putative target of SR45 splicing activity (SAR) but was also identified in a set of 918 differentially expressed genes in *sr45-1* seedlings (Xing *et al*., 2015) and as down-regulated in roots of adult *sr45-1* plants here. Even though several phenotypes, including reduced growth rate of the primary root (Pirone *et al*., 2017), of the *pwd/gwd2* mutant are a surprising reminiscence of the root and reproductive defects we observed in *sr45-1* **(Figure 2** and **Figure 7)**, it was recently shown that the down-regulation of *PWD/GWD2* expression does not correlate with a protein decrease in the *sr45-1* mutant inflorescences (Chen *et al*., 2019). However, because the PWD/GWD2 function is restricted to the companion cells (Glaring *et al*., 2007) and consists in sustaining the phloem loading of polysaccharides in source organs (Pirone *et al*., 2017), the absence of protein accumulation changes in inflorescences is not relevant in regards to the primary root cellular function played by this protein (Pirone *et al*., 2017). Therefore, the reproductive defects of *sr45-1* might not be solely associated to the down-regulation of *PWD/GWD2*, which is confirmed by the fact that Fe supplementation is able to rescue the reproductive defects of the mutant **(Figure 8)** in the absence of regulation of *PWD/GWD2* **(Supplemental Figure S11)**. These results strongly suggest that part of the silique and seed developmental defects is related to altered metal homeostasis in reproductive organs, which is known to tremendously affect anther dehiscence, pollen development, fruit development, seed quality and seed yield (Verbruggen and Hermans, 2013; Guo *et al*., 2016; Singh and Reddy, 2017).

Supporting this hypothesis, *sr45-1* mutant seeds contain decreased amounts of Mn as well as increased levels of macronutrients K, Mg and Zn, possibly accounting for the reproductive defects upon water irrigation, i.e. in the absence of Fe supplementation **(Figure 7)**. Previously, the expression of several ion transporter-encoding genes was shown to be deregulated in *sr45-1* inflorescences (Zhang *et al*., 2017). Indeed, *AtHMP37* (*AT4G27590*) and *AtHMP27* (*AT3G24450*), *PCR2* and *HIPP22*, respectively encoding two heavy metal transport/detoxification superfamily proteins (Li *et al*., 2020), a zinc transporter (Song *et al*., 2010) and a cadmium-detoxifying protein (Tehseen *et al*., 2010), are all up-regulated in *sr45-1* inflorescences (Zhang *et al*., 2017). The member of the heavy metal transport/detoxification superfamily, *AtHMP37*, is a SDR gene (in both roots and inflorescences) identified as a putative RNA target of SR45 (SAR) **(Supplemental Table S6)** (Xing *et al*., 2015; Zhang *et al*., 2017) that is up-regulated in *sr45-1* roots **(Supplemental Figure S13)**. On the contrary, the *NFP7.1* gene, which encodes a nitrate transporter (Babst *et al*., 2019), and the NRT1.7 nitrate transporter (Fan *et al*., 2009; Liu *et al*., 2017b), were shown to be down-regulated in *sr45-1* inflorescences (Chen *et al*., 2019). It is well known that the N status affects Mg and Fe homeostasis (e.g. *FIT* regulation) (García *et al*., 2010; Curie and Mari, 2017; Liu *et al*., 2017a; Kailasam *et al*., 2018; Liu *et al*., 2018) and that a crosstalk exists between Zn and Fe homeostasis through at least IRT1, ZIF1 and FRD3 (Haydon *et al*., 2012; Charlier *et al*., 2015; Scheepers *et al*., 2020; Hanikenne *et al*., 2021). It is therefore possible that changes in the levels of micro- and macronutrients in seeds are the result of deregulation in the expression of nutrient transporters or *vice versa*, the causality between these observations being difficult to determine from our data. In fact, the mutant root growth is altered by Fe supply from the first week of development (see **Supplemental Figure S12)** and continues to be drastically affected upon extensive exposure **(Figure 2)**.

Surprisingly, while Fe concentration was diminished in the *sr45-1* mutant shoots upon both Fe deficiency and sufficiency **(Supplemental Figure S2)**, it is very similar in seeds of Col-0 and *sr45-1* **(Supplemental Figure S10D)**. This suggest that the cause of the *sr45-1* developmental defects in reproductive organs is not Fe starvation of the embryo but rather a consequence of intricate ionome perturbations. In fact, Zn concentration in seeds obtained from plants watered with solely tap water was higher (84.5 %) in the mutant compared to the WT, suggesting that the radicle local environment contained a higher Zn pool, which may result in a local Fe deficiency as previously observed (Dučić and Polle, 2007; Lei *et al*., 2007; Kawachi *et al*., 2009; Fukao *et al*., 2011; Shanmugam *et al*., 2011; Marschner and Marschner, 2012; Millaleo *et al*., 2013; Scheepers *et al*., 2020). The constitutive induction of the Fe uptake machinery **(Figure 5)** in the *sr45-1* seedlings would reflect this ionome and possibly Fe defect in the seeds. However, the primary root growth of seeds collected from *sr45-1* plants supplemented with Fe was not significantly improved compared to seeds collected from plants watered with tap water **(Supplemental Figure S12)**. It is therefore still unclear how altered partitioning of metals in the mutant seeds affects the primary growth of the seedling.

It is however evident that *sr45-1* seedlings and plants exhibit several alterations of metal homeostasis: (i) Fe and H_2_O_2_ accumulation within the root vasculature; (ii) Mn and Zn accumulation in roots and (iii) reduced shoot metal (Fe, Zn, Mn) concentrations, resulting in reduced biomass and chlorosis upon changes in Fe supply. RNA-Seq and qRT-PCR analyses suggest that mutant roots displayed a basal induction of Fe-starvation-responsive genes in control conditions with less drastic expression changes compared to Col-0 upon Fe deficiency (1350 *vs* 2442 DEGs) in adult plants **(Figure 4, Supplemental Figures S7 and S13)** and a stronger response in seedlings (**Figure 5, Supplemental Figure S6**).

Our working hypothesis therefore stands in favor of a scenario where the defective seed ionome affects the development of the embryo, which consequently results in an induction of the Fe uptake machinery in seedlings upon germination **(Figure 5A-D)** in an attempt to compensate for the metal starvation **(Figure 7J)**. The down-regulation of *FRD3* expression in seedlings **(Figure 5E)** would reduce root-to-shoot translocation of citrate-complexed metals that may reveal toxic when in excess in aerial parts (i.e. Fe, Zn). This ultimately results in lower shoot biomass and metal concentrations **(Figure 1 and Supplemental Figure S2)**, together with Fe and Zn toxicity in roots **(Figure 3)**, including the accumulation of H_2_O_2_ **(Figure 1F** and **Supplemental Figure S5B)**. If ionome and growth perturbations persist in mutant adult plants **(Figure 7G-J)**, their effect on Fe and metal homeostasis gene expression appear mitigated **(Figure 4** and **Figure 5)**.

## Conclusion

We reported here that some of the *sr45-1* mutant phenotype result from a severe alteration in metal mobilization, localization and transport. We also showed that exogenous application of Fe can rescue the reproductive defects of the mutant. Our data fine-tune our understanding of the physiological responses impaired in the mutant, such as the oxidative response, metal homeostasis and the phenylpropanoid pathway.

## Methods

### Plant material and growth conditions

All experiments were conducted under a 8-h-light (100 µmol m^-2^ s^-1^)/16-h-dark regime in a climate-controlled growth chamber (21°C). *Arabidopsis thaliana* ecotype Columbia-0 (Col-0) was used as wild-type, and seeds from *sr45-1* mutant (SALK_004132, Col-0 background) and *frd3-7* mutant (SALK_122235, Col-0 background) were obtained from the SALK collection. Seeds were surface-sterilized and germinated on 1/2 Murashige and Skoog (MS) medium (Duchefa Biochimie) supplemented with sucrose (1% w/v, Duchefa Biochimie) and Select Agar (0.8% w/v, Sigma-Aldrich) and stratified in the dark at 4°C for 48h. For hydroponics experiments, 3-week-old seedlings were transferred in hydroponic trays (Araponics) and cultivated for 3 weeks in control Hoagland medium, followed by 3 weeks of experimental conditions. The nutrient medium was renewed with fresh solution once a week and 3 days prior to the harvesting. The control Hoagland medium included 10 µM FeIII-HBED [N,N’-di(2-hydroxybenzyl) ethylenediamine N,N’-diacetic acid monohydrochloride], 1 µM Zn (ZnSO_4_.H_2_O) and 5 µM Mn (MnSO_4_.H_2_O) as reported in (Talke *et al*., 2006; Hanikenne *et al*., 2008; Scheepers *et al*., 2020). Fe, Zn and Mn were respectively added or omitted from medium as described.

Unless stated otherwise, for root length measurement (including treatments with 120 µM of DMF-solubilized fraxetin), Perls and hydrogen peroxide staining, acidification capacity assay, FeIII chelate reductase activity assay and citrate content measurement, surface-sterilized seeds were directly sown on square plastic Petri plates (Greiner Bio-One) containing modified Hoagland supplemented with sucrose (1% w/v, Duchefa Biochimie), FeIII-HBED (0 µM to 50 µM) and agar (0.8% w/v, Agar Type M, Sigma-Aldrich), and grown vertically after stratification.

### Seed number per silique, seed morphology, silique size and roots length

3-weeks-old seedlings were transferred in soil and daily watered with water (referred to as –Fe). In addition of this treatment, half of the seedlings from each genotype were weekly supplied with a sequestrene solution (referred to as +Fe, 0.1 g/L, Liro N.V.) until the completion of silique development and seed maturation. Siliques were then harvested before dehiscence, and their size was determined using the segmented line tool on ImageJ. After an incubation of two weeks in 95% ethanol at room temperature, the seed number per silique was determined under a Nikon SMZ1500 stereomicroscope. Seeds were finally photographed using a Nikon SMZ1500 stereomicroscope equipped with a Nikon Digital Sight DS-5M camera in order to determine seed length and width.

### Perls and hydrogen peroxide staining

For Perls staining, roots of *Arabidopsis thaliana* Col-0 and *sr45-1* mutant seedlings (or 9-week-old plants) were vacuum infiltrated with HCl (4%, v/v) and K-Ferrocyanide (II) (4%, w/v, Sigma-Aldrich) (1/1) for 15 minutes. The reaction was continued for an additional 30 minutes at room temperature and was then stopped by substituting the solution by distilled water (Roschzttardtz *et al*., 2009). Observation was realized using a Nikon SMZ1500 stereomicroscope equipped with a Nikon Digital Sight DS-5M camera.

Hydrogen peroxide (H_2_O_2_) was detected according to (Baliardini *et al*., 2015). Roots samples were vacuum infiltrated with 3-3’-diaminobenzidine tetrahydrochloride (1.25 mg/mL, DAB, Sigma-Aldrich), Tween-20 (0.05% v/v) and 200 mM Na_2_HPO_4_, then incubated in the same solution for a total of one hour. They were subsequently bleached in acetic-acid/glycerol/ethanol (1/1/3) during 5 minutes at 100°C, and stored in glycerol/ethanol (1/4) before further analysis. Samples were observed under a stereomicroscope as above.

### Acidification capacity and Ferric (FeIII) chelate reductase activity assays

For the measure of acidification capacity, roots from a pool of 5 seedlings were incubated in 0.005% bromocresol purple (Roth) during 24 hours in the dark. *A_433_* of the protonated form of the dye was then measured and expressed relative to the root weight of sample (Santi and Schmidt, 2009; El-Ashgar *et al*., 2012).

The ferric chelate reductase (FCR) activity was measured on roots from a pool of 5 seedlings. Samples were immerged in a reductase solution containing FeIII-EDTA (0.1 mM, Roth) and FerroZine (0.3 mM, Acros Organics) for 30 minutes in the dark. *A_562_* of the FeII-FerroZine complex was then determined. The final calculation included the root weight of sample and the molar extinction coefficient of the complex (28.6 mM^-1^.cm^-1^) (Yi and Guerinot, 1996).

### Citrate content

Citrate determination was performed as previously described (Schvartzman *et al*., 2018). Briefly, 100 mg of roots were frozen and grinded in liquid nitrogen with a mortar and pestle. Samples were resuspended in 1 mL of distilled water, and the pH was adjusted to 7 – 8 with 1 M KOH. Samples were deproteinized using 100 µL of 1 M perchloric acid, and citrate content was measured using a citric acid assay kit according to the manufacturer’s protocol (BioSentec, France).

### Total chlorophyll and carotenoid content

Total chlorophyll content was determined from three to six young seedlings or the rosette of one 9-week-old plant. Plants were weighed, then incubated in the dark during 72h or seven days in 95% ethanol. Discolored plants were removed and the solution was submitted to spectroscopic analysis. The total amount of chlorophyll was calculated using the equation: 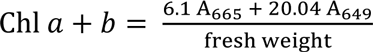 (Wintermans and de Mots, 1965). Carotenoid content (xanthophylls and carotenes) was determined for the same samples using the equation: 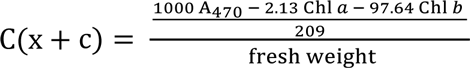 (Lichtenthaler and Buschmann, 2001).

### Gene expression analysis

Total RNAs were extracted from 100 mg of plant tissues (entire plants or roots) using NucleoSpin RNA Plant kit (Macherey Nagel) as per manufacturer’s instruction. cDNAs were synthesized from 1 µg of total RNAs using oligo(dT) and the RevertAid H Minus First Strand cDNA Synthesis Kit (Fisher Scientific). Quantitative PCR reactions were performed in a QuantStudio5 (Applied Biosystems) using 384-well plates and Takyon Low Rox SYBR MasterMix dTTP Blue (Eurogentec) on material from three independent biological experiments, and a total of three technical repeats were run for each combination of cDNA and primer pair. Gene expression was normalized relative to At1g58050 as described (Pfaffl, 2001). *At1g58050* expression was the more stable among all tested references (*EF1α* and *UBQ10*) (Czechowski *et al*., 2005; Spielmann *et al*., 2020). **Supplemental Table S7** shows the primers used for these experiments.

Upon harvesting, plant root tissues were blotted dry, immediately frozen in liquid nitrogen and stored at −80°C. Total RNAs were prepared using 100 mg of homogenized tissues and RNeasy Plant Mini kit with on-column DNase treatment (Qiagen). Libraries for RNA-Seq were prepared from 1 µg of total RNAs with the TruSeq Stranded mRNA Library Prep Kit (Illumina, CA, USA), multiplexed and sequenced in two runs with an Illumina NextSeq500 device (high throughput mode, 75 base single-end reads) at the GIGA-R Sequencing platform (University of Liège), yielding on average ∼22 million reads per sample. Read quality was assessed using FastQC (v0.10.1, http://www.bioinformatics.babraham.ac.uk/projects/fastqc/). Quality trimming and removal of adapters were conducted using Trimmomatic (v0.32, Bolger *et al*., 2014), with the following parameters: trim bases with quality score lower than Q26 in 5’ and 3’ of reads; remove any reads with Q<26 in any sliding window of 10 bases; crop 1 base in 3’ of all reads, and discard reads shorter than 70 bases. Overall quality filtering discarded between 7 and 9% of the raw reads. The Arabidopsis reference genome sequence (TAIR10) and annotation (201606 version) files were downloaded from Araport on Sept 16, 2016 (www.araport.org). Read mapping on the genome was achieved using TopHat (v2.1.1), with the following parameters: -- read-mismatches 2; --min-intron-length 40; --max-intron-length 2000; 2 --report-secondary-alignments; --no-novel-juncs and providing an indexed genome annotation file. Raw read counts were obtained using htseq-count (v0.6.1p1) and differentially expressed genes were identified by pairwise comparisons with the DESeq2 package (v1.12.3, Love *et al*., 2014). Genes were retained as differentially expressed when the log_2_ fold-change (FC) was > 1 or < −1, with a false discovery rate (FDR, Benjamini-Hochberg) adjusted *p-value* of < 0.05. Principal Component Analysis plots (PCA) were created with the *PlotPCA* function from R using rlog transformed data (Beginner’s guide, DESeq2 package, May 13, 2014, http://www.bioconductor.org/packages/2.14/bioc/vignettes/DESeq2/inst/doc/beginner.pdf). GO enrichment analyses were conducted using the Thalemine tool on Araport (http://www.araport.org). The heatmaps were constructed using the heatmap.2 function of the gplots R package.

### Analysis of metal content

Seed (–Fe and +Fe), root and shoot tissues of wild-type plants (Col-0) and *sr45-1* mutant plants were harvested separately. Root tissues were desorbed and washed as previously described (Talke *et al*., 2006), and seed and shoot tissues were rinsed in distillated water. 1 to 30 mg of dried tissue were digested and prepared as described (Nouet *et al*., 2015). Metal content was determined by ICP-AES (inductively coupled plasma-atomic emission spectroscopy) (Vista AX, Varian).

### Statistical analysis

All data evaluation and statistics were done using GraphPad Prism 7 (GraphPad Software v7.00).

## Accession numbers and information

All gene sequences are available through *The Arabidopsis Information Resource* (TAIR, http://www.arabidopsis.org/), with this accession number: *Arabidopsis SR45* (AT1G16610), *AHA2* (AT4G30190), *bHLH100* (AT2G41240), *bHLH101* (AT5G04150), *bHLH11* (AT4G36060), *bHLH18* (AT2G22750), *bHLH19* (AT2G22760), *bHLH20* (AT2G22770), *bHLH25* (AT4G37850), *bHLH38* (AT3G56970), *bHLH39* (AT3G56980), *FIT* (AT2G28160), *FRD3* (AT3G08040), *FRO2* (AT1G01580), *IRT1* (AT4G19690), *MYB15* (AT3G23250), *MYB72* (AT1G56160), *NAS1* (AT5G04950), *NAS2* (AT5G56080), *NAS3* (AT1G09240), *NAS4* (AT1G56430), and *S8H* (AT3G12900). The *Arabidopsis thaliana sr45-1* T-DNA insertion (SALK_004132) and *frd3-7* T-DNA insertion (SALK_122235) lines were available at the SALK collection. The RNA-Seq reads have been deposited in the National Center for Biotechnology Information (NCBI) Sequence Read Archive Database with BioProject identification number [XXX].

## Supplemental data

**Supplemental Figure S1.** Number of potential SR45 targets showing iron deficiency responsiveness.

**Supplemental Figure S2.** Micronutrient concentrations in *sr45-1* shoots.

**Supplemental Figure S3.** Macronutrient concentrations in *sr45-1* tissues.

**Supplemental Figure S4.** Phenotype characterization of wild-type and *sr45-1* seedlings and plants upon iron deficiency and iron excess.

**Supplemental Figure S5.** Toxicity of manganese and zinc in roots and shoots.

**Supplemental Figure S6.** Fe uptake machinery activity in *sr45-1* seedlings.

**Supplemental Figure S7.** Regulation of FIT-interacting coactivators in *sr45-1* adult plants.

**Supplemental Figure S8.** Transcriptomic analysis in roots upon iron deficiency and sufficiency.

**Supplemental Figure S9.** Effect of exogenous application of coumarins on *sr45-1* shoots.

**Supplemental Figure S10.** Effect of irrigation on stem height and flowering.

**Supplemental Figure S11.** SR45 differentially regulated genes (SDRs) involved in reproductive development.

**Supplemental Figure S12.** Root length of *sr45-1* seedlings upon iron deficiency and iron excess.

**Supplemental Figure S13.** Transcriptomic analysis in roots upon iron deficiency and sufficiency.

**Supplemental Table S1.** Significantly enriched categories among SR45-associated RNAs.

**Supplemental Table S2.** Lists of differentially expressed genes from all pairwise comparisons described in Figure 4.

**Supplemental Table S3.** Significantly enriched categories among DEGs identified in roots of WT and *sr45-1* upon iron deficiency and sufficiency.

**Supplemental Table S4.** Col-0 differentially regulated genes (CDR) and SR45 differentially regulated genes (SDR) from all pairwise comparisons described in Figure 4.

**Supplemental Table S5.** Significantly enriched categories among CDR genes and SDR genes.

**Supplemental Table S6.** Comparison of SDR genes identified in roots with SR45-associated RNAs and SDR genes identified in inflorescences.

**Supplemental Table S7.** List of primers used in this study.

## Acknowledgments

We thank Dr. Julien Spielmann for helpful discussions, and Dr. S. Thiriet-Rupert and A. Bertrand for their help with RStudio. We thank Prof. M. Carnol, A. Degueldre and B. Bosman for their help with ICP-AES analyses. Funding was provided by the “Fonds de la Recherche Scientifique–FNRS” (FRFC-1.E049.15, PDR-T.0206.13, CDR-J.0009.17, PDR-T0120.18, CDR-J.0082.21). M.H. is Senior Research Associate of the F.R.S.-FNRS. S.F. was a doctoral fellow (F.R.I.A.). No conflict of interest declared.

## Literature cited

Albaqami, M., Laluk, K., and Reddy, A.S.N. (2019). The Arabidopsis splicing regulator SR45 confers salt tolerance in a splice isoform-dependent manner. Plant Mol Biol 100, 379–390.

Ali, G.S., Palusa, S.G., Golovkin, M., Prasad, J., Manley, J.L., and Reddy, A.S. (2007). Regulation of plant developmental processes by a novel splicing factor. PLoS One 2, e471.

Babst, B.A., Gao, F., Acosta-Gamboa, L.M., Karve, A., Schueller, M.J., and Lorence, A. (2019). Three NPF genes in Arabidopsis are necessary for normal nitrogen cycling under low nitrogen stress. Plant Physiol Biochem 143, 1–10.

Baliardini, C., Meyer, C.L., Salis, P., Saumitou-Laprade, P., and Verbruggen, N. (2015). CATION EXCHANGER1 Cosegregates with Cadmium Tolerance in the Metal Hyperaccumulator Arabidopsis halleri and Plays a Role in Limiting Oxidative Stress in Arabidopsis Spp. Plant Physiol 169, 549–559.

Barberon, M., Vermeer, J.E., De Bellis, D., Wang, P., Naseer, S., Andersen, T.G., Humbel, B.M., Nawrath, C., Takano, J., Salt, D.E., and Geldner, N. (2016). Adaptation of Root Function by Nutrient-Induced Plasticity of Endodermal Differentiation. Cell 164, 447–459.

Barta, A., Kalyna, M., and Reddy, A.S. (2010). Implementing a rational and consistent nomenclature for serine/arginine-rich protein splicing factors (SR proteins) in plants. Plant Cell 22, 2926–2929.

Baxter, I., Hosmani, P.S., Rus, A., Lahner, B., Borevitz, J.O., Muthukumar, B., Mickelbart, M.V., Schreiber, L., Franke, R.B., and Salt, D.E. (2009). Root suberin forms an extracellular barrier that affects water relations and mineral nutrition in Arabidopsis. PLoS Genet 5, e1000492.

Bolger, A.M., Lohse, M., and Usadel, B. (2014). Trimmomatic: a flexible trimmer for Illumina sequence data. Bioinformatics 30, 2114–2120.

Califice, S., Baurain, D., Hanikenne, M., and Motte, P. (2012). A single ancient origin for prototypical serine/arginine-rich splicing factors. Plant Physiol 158, 546–560.

Carvalho, R.F., Carvalho, S.D., and Duque, P. (2010). The plant-specific SR45 protein negatively regulates glucose and ABA signaling during early seedling development in Arabidopsis. Plant Physiol 154, 772–783.

Carvalho, R.F., Szakonyi, D., Simpson, C.G., Barbosa, I.C., Brown, J.W., Baena-González, E., and Duque, P. (2016). The Arabidopsis SR45 Splicing Factor, a Negative Regulator of Sugar Signaling, Modulates SNF1-Related Protein Kinase 1 Stability. Plant Cell 28, 1910–1925.

Castaings, L., Caquot, A., Loubet, S., and Curie, C. (2016). The high-affinity metal Transporters NRAMP1 and IRT1 Team up to Take up Iron under Sufficient Metal Provision. Sci Rep 6, 37222.

Chamala, S., Feng, G., Chavarro, C., and Barbazuk, W.B. (2015). Genome-wide identification of evolutionarily conserved alternative splicing events in flowering plants. Front Bioeng Biotechnol 3, 33.

Charlier, J.B., Polese, C., Nouet, C., Carnol, M., Bosman, B., Krämer, U., Motte, P., and Hanikenne, M. (2015). Zinc triggers a complex transcriptional and post-transcriptional regulation of the metal homeostasis gene FRD3 in Arabidopsis relatives. J Exp Bot 66, 3865–3878.

Chen, S.L., Rooney, T.J., Hu, A.R., Beard, H.S., Garrett, W.M., Mangalath, L.M., Powers, J.J., Cooper, B., and Zhang, X.N. (2019). Quantitative Proteomics Reveals a Role for SERINE/ARGININE-Rich 45 in Regulating RNA Metabolism and Modulating Transcriptional Suppression via the ASAP Complex in Arabidopsis thaliana. Front Plant Sci 10, 1116.

Chen, T., Cui, P., Chen, H., Ali, S., Zhang, S., and Xiong, L. (2013). A KH-domain RNA-binding protein interacts with FIERY2/CTD phosphatase-like 1 and splicing factors and is important for pre-mRNA splicing in Arabidopsis. PLoS Genet 9, e1003875.

Chezem, W.R., Memon, A., Li, F.S., Weng, J.K., and Clay, N.K. (2017). SG2-Type R2R3-MYB Transcription Factor MYB15 Controls Defense-Induced Lignification and Basal Immunity in Arabidopsis. Plant Cell 29, 1907–1926.

Colangelo, E.P., and Guerinot, M.L. (2004). The essential basic helix-loop-helix protein FIT1 is required for the iron deficiency response. Plant Cell 16, 3400–3412.

Cui, Y., Chen, C.L., Cui, M., Zhou, W.J., Wu, H.L., and Ling, H.Q. (2018). Four IVa bHLH Transcription Factors Are Novel Interactors of FIT and Mediate JA Inhibition of Iron Uptake in Arabidopsis. Mol Plant 11, 1166–1183.

Curie, C., and Mari, S. (2017). New routes for plant iron mining. New Phytol 214, 521–525.

Czechowski, T., Stitt, M., Altmann, T., Udvardi, M.K., and Scheible, W.R. (2005). Genome-wide identification and testing of superior reference genes for transcript normalization in Arabidopsis. Plant Physiol 139, 5–17.

Dong, C., He, F., Berkowitz, O., Liu, J., Cao, P., Tang, M., Shi, H., Wang, W., Li, Q., Shen, Z., Whelan, J., and Zheng, L. (2018). Alternative Splicing Plays a Critical Role in Maintaining Mineral Nutrient Homeostasis in Rice (Oryza sativa). Plant Cell 30, 2267–2285.

Dučić, T., and Polle, A. (2007). Manganese toxicity in two varieties of Douglas fir (Pseudotsuga menziesii var. viridis and glauca) seedlings as affected by phosphorus supply. Functional Plant Biology 34, 31–40.

Durrett, T.P., Gassmann, W., and Rogers, E.E. (2007). The FRD3-mediated efflux of citrate into the root vasculature is necessary for efficient iron translocation. Plant Physiol 144, 197–205.

El-Ashgar, N.M., El-Basioni, A., El-Nahhal, I.M., Zourob, S.M., El-Agez, T.M., and Taya, S.A. (2012). Sol-Gel Thin Films Immobilized with Bromocresol Purple pH-Sensitive Indicator in Presence of Surfactants. ISRN Analytical Chemistry 2012.

Fan, S.C., Lin, C.S., Hsu, P.K., Lin, S.H., and Tsay, Y.F. (2009). The Arabidopsis nitrate transporter NRT1.7, expressed in phloem, is responsible for source-to-sink remobilization of nitrate. Plant Cell 21, 2750–2761.

Fourcroy, P., Sisó-Terraza, P., Sudre, D., Saviron, M., Reyt, G., Gaymard, F., Abadía, A., Abadia, J., Álvarez-Fernández, A., and Briat, J.F. (2014). Involvement of the ABCG37 transporter in secretion of scopoletin and derivatives by Arabidopsis roots in response to iron deficiency. New Phytol 201, 155–167.

Fukao, Y., Ferjani, A., Tomioka, R., Nagasaki, N., Kurata, R., Nishimori, Y., Fujiwara, M., and Maeshima, M. (2011). iTRAQ analysis reveals mechanisms of growth defects due to excess zinc in Arabidopsis. Plant Physiol 155, 1893–1907.

García, M.J., Lucena, C., Romera, F.J., Alcántara, E., and Pérez-Vicente, R. (2010). Ethylene and nitric oxide involvement in the up-regulation of key genes related to iron acquisition and homeostasis in Arabidopsis. J Exp Bot 61, 3885–3899.

Geldner, N. (2013). The endodermis. Annu Rev Plant Biol 64, 531–558.

Glaring, M.A., Zygadlo, A., Thorneycroft, D., Schulz, A., Smith, S.M., Blennow, A., and Baunsgaard, L. (2007). An extra-plastidial alpha-glucan, water dikinase from Arabidopsis phosphorylates amylopectin in vitro and is not necessary for transient starch degradation. J Exp Bot 58, 3949–3960.

Golovkin, M., and Reddy, A.S. (1999). An SC35-like protein and a novel serine/arginine-rich protein interact with Arabidopsis U1-70K protein. J Biol Chem 274, 36428–36438.

Green, L.S., and Rogers, E.E. (2004). FRD3 controls iron localization in Arabidopsis. Plant Physiol 136, 2523–2531.

Guo, W., Nazim, H., Liang, Z., and Yang, D. (2016). Magnesium deficiency in plants: An urgent problem. The Crop Journal 4, 83–91.

Hanikenne, M., Esteves, S.M., Fanara, S., and Rouached, H. (2021). Coordinated homeostasis of essential mineral nutrients: a focus on iron. J Exp Bot 72, 2136–2153.

Hanikenne, M., Talke, I.N., Haydon, M.J., Lanz, C., Nolte, A., Motte, P., Kroymann, J., Weigel, D., and Krämer, U. (2008). Evolution of metal hyperaccumulation required cis-regulatory changes and triplication of HMA4. Nature 453, 391–395.

Haydon, M.J., Kawachi, M., Wirtz, M., Hillmer, S., Hell, R., and Krämer, U. (2012). Vacuolar nicotianamine has critical and distinct roles under iron deficiency and for zinc sequestration in Arabidopsis. Plant Cell 24, 724–737.

Hindt, M.N., Akmakjian, G.Z., Pivarski, K.L., Punshon, T., Baxter, I., Salt, D.E., and Guerinot, M.L. (2017). BRUTUS and its paralogs, BTS LIKE1 and BTS LIKE2, encode important negative regulators of the iron deficiency response in Arabidopsis thaliana. Metallomics 9, 876–890.

Ivanov, R., Brumbarova, T., and Bauer, P. (2012). Fitting into the harsh reality: regulation of iron-deficiency responses in dicotyledonous plants. Mol Plant 5, 27–42.

Jakoby, M., Wang, H.Y., Reidt, W., Weisshaar, B., and Bauer, P. (2004). FRU (BHLH029) is required for induction of iron mobilization genes in Arabidopsis thaliana. FEBS Lett 577, 528–534.

Jeong, S. (2017). SR Proteins: Binders, Regulators, and Connectors of RNA. Mol Cells 40, 1–9.

Kailasam, S., Wang, Y., Lo, J.C., Chang, H.F., and Yeh, K.C. (2018). S-Nitrosoglutathione works downstream of nitric oxide to mediate iron-deficiency signaling in Arabidopsis. Plant J 94, 157–168.

Kamiya, T., Borghi, M., Wang, P., Danku, J.M., Kalmbach, L., Hosmani, P.S., Naseer, S., Fujiwara, T., Geldner, N., and Salt, D.E. (2015). The MYB36 transcription factor orchestrates Casparian strip formation. Proc Natl Acad Sci U S A 112, 10533–10538.

Kawachi, M., Kobae, Y., Mori, H., Tomioka, R., Lee, Y., and Maeshima, M. (2009). A mutant strain Arabidopsis thaliana that lacks vacuolar membrane zinc transporter MTP1 revealed the latent tolerance to excessive zinc. Plant Cell Physiol 50, 1156–1170.

Kobayashi, T., Nagasaka, S., Senoura, T., Itai, R.N., Nakanishi, H., and Nishizawa, N.K. (2013). Iron-binding haemerythrin RING ubiquitin ligases regulate plant iron responses and accumulation. Nat Commun 4, 2792.

Laloum, T., Martín, G., and Duque, P. (2018). Alternative Splicing Control of Abiotic Stress Responses. Trends Plant Sci 23, 140–150.

Le, C.T., Brumbarova, T., Ivanov, R., Stoof, C., Weber, E., Mohrbacher, J., Fink-Straube, C., and Bauer, P. (2016). ZINC FINGER OF ARABIDOPSIS THALIANA12 (ZAT12) Interacts with FER-LIKE IRON DEFICIENCY-INDUCED TRANSCRIPTION FACTOR (FIT) Linking Iron Deficiency and Oxidative Stress Responses. Plant Physiol 170, 540–557.

Lei, Y., Korpelainen, H., and Li, C. (2007). Physiological and biochemical responses to high Mn concentrations in two contrasting Populus cathayana populations. Chemosphere 68, 686–694.

Li, J., Zhang, M., Sun, J., Mao, X., Wang, J., Liu, H., Zheng, H., Li, X., Zhao, H., and Zou, D. (2020). Heavy Metal Stress-Associated Proteins in Rice and Arabidopsis: Genome-Wide Identification, Phylogenetics, Duplication, and Expression Profiles Analysis. Front Genet 11, 477.

Lichtenthaler, H.K., and Buschmann, C. (2001). Chlorophylls and Carotenoids: Measurement and Characterization by UV-VIS Spectroscopy. Current Protocols in Food Analytical Chemistry F4, 1–F4.3.8.

Lingam, S., Mohrbacher, J., Brumbarova, T., Potuschak, T., Fink-Straube, C., Blondet, E., Genschik, P., and Bauer, P. (2011). Interaction between the bHLH transcription factor FIT and ETHYLENE INSENSITIVE3/ETHYLENE INSENSITIVE3-LIKE1 reveals molecular linkage between the regulation of iron acquisition and ethylene signaling in Arabidopsis. Plant Cell 23, 1815–1829.

Liu, M., Liu, X.X., He, X.L., Liu, L.J., Wu, H., Tang, C.X., Zhang, Y.S., and Jin, C.W. (2017a). Ethylene and nitric oxide interact to regulate the magnesium deficiency-induced root hair development in Arabidopsis. New Phytol 213, 1242–1256.

Liu, M., Zhang, H., Fang, X., Zhang, Y., and Jin, C. (2018). Auxin Acts Downstream of Ethylene and Nitric Oxide to Regulate Magnesium Deficiency-Induced Root Hair Development in Arabidopsis thaliana. Plant Cell Physiol 59, 1452–1465.

Liu, W., Sun, Q., Wang, K., Du, Q., and Li, W.X. (2017b). Nitrogen Limitation Adaptation (NLA) is involved in source-to-sink remobilization of nitrate by mediating the degradation of NRT1.7 in Arabidopsis. New Phytol 214, 734–744.

Long, T.A., Tsukagoshi, H., Busch, W., Lahner, B., Salt, D.E., and Benfey, P.N. (2010). The bHLH transcription factor POPEYE regulates response to iron deficiency in Arabidopsis roots. Plant Cell 22, 2219–2236.

Love, M.I., Huber, W., and Anders, S. (2014). Moderated estimation of fold change and dispersion for RNA-seq data with DESeq2. Genome Biology 15, 550.

Manley, J.L., and Krainer, A.R. (2010). A rational nomenclature for serine/arginine-rich protein splicing factors (SR proteins). Genes Dev 24, 1073–1074.

Marschner, H., and Marschner, P. (2012). Marschner’s Mineral Nutrition of Higher Plants (3rd edition). Academic Press.

Meyer, K., Koester, T., and Staiger, D. (2015). Pre-mRNA Splicing in Plants: In Vivo Functions of RNA-Binding Proteins Implicated in the Splicing Process. Biomolecules 5, 1717–1740.

Millaleo, R., Reyes-Díaz, M., Alberdi, M., Ivanov, A.G., Krol, M., and Hüner, N.P. (2013). Excess manganese differentially inhibits photosystem I versus II in Arabidopsis thaliana. J Exp Bot 64, 343–354.

Mladenka, P., Macáková, K., Zatloukalová, L., Reháková, Z., Singh, B.K., Prasad, A.K., Parmar, V.S., Jahodár, L., Hrdina, R., and Saso, L. (2010). In vitro interactions of coumarins with iron. Biochimie 92, 1108–1114.

Morrissey, J., Baxter, I.R., Lee, J., Li, L., Lahner, B., Grotz, N., Kaplan, J., Salt, D.E., and Guerinot, M.L. (2009). The ferroportin metal efflux proteins function in iron and cobalt homeostasis in Arabidopsis. Plant Cell 21, 3326–3338.

Nouet, C., Charlier, J.B., Carnol, M., Bosman, B., Farnir, F., Motte, P., and Hanikenne, M. (2015). Functional analysis of the three HMA4 copies of the metal hyperaccumulator Arabidopsis halleri. J Exp Bot 66, 5783–5795.

Nouet, C., Motte, P., and Hanikenne, M. (2011). Chloroplastic and mitochondrial metal homeostasis. Trends Plant Sci 16, 395–404.

Palmer, C.M., Hindt, M.N., Schmidt, H., Clemens, S., and Guerinot, M.L. (2013). MYB10 and MYB72 are required for growth under iron-limiting conditions. PLoS Genet 9, e1003953.

Palusa, S.G., Ali, G.S., and Reddy, A.S. (2007). Alternative splicing of pre-mRNAs of Arabidopsis serine/arginine-rich proteins: regulation by hormones and stresses. Plant J 49, 1091–1107.

Palusa, S.G., and Reddy, A.S. (2010). Extensive coupling of alternative splicing of pre-mRNAs of serine/arginine (SR) genes with nonsense-mediated decay. New Phytol 185, 83–89.

Petridis, A., Döll, S., Nichelmann, L., Bilger, W., and Mock, H.P. (2016). Arabidopsis thaliana G2-LIKE FLAVONOID REGULATOR and BRASSINOSTEROID ENHANCED EXPRESSION1 are low-temperature regulators of flavonoid accumulation. New Phytol 211, 912–925.

Pfaffl, M.W. (2001). A new mathematical model for relative quantification in real-time RT-PCR. Nucleic Acids Res 29, e45.

Pirone, C., Gurrieri, L., Gaiba, I., Adamiano, A., Valle, F., Trost, P., and Sparla, F. (2017). The analysis of the different functions of starch-phosphorylating enzymes during the development of Arabidopsis thaliana plants discloses an unexpected role for the cytosolic isoform GWD2. Physiol Plant 160, 447–457.

Rajniak, J., Giehl, R.F.H., Chang, E., Murgia, I., von Wirén, N., and Sattely, E.S. (2018). Biosynthesis of redox-active metabolites in response to iron deficiency in plants. Nat Chem Biol 14, 442–450.

Ravet, K., Touraine, B., Boucherez, J., Briat, J.F., Gaymard, F., and Cellier, F. (2009). Ferritins control interaction between iron homeostasis and oxidative stress in Arabidopsis. Plant J 57, 400–412.

Remy, E., Cabrito, T.R., Batista, R.A., Hussein, M.A., Teixeira, M.C., Athanasiadis, A., Sá-Correia, I., and Duque, P. (2014). Intron retention in the 5’UTR of the novel ZIF2 transporter enhances translation to promote zinc tolerance in Arabidopsis. PLoS Genet 10, e1004375.

Reyt, G., Boudouf, S., Boucherez, J., Gaymard, F., and Briat, J.F. (2015). Iron- and ferritin-dependent reactive oxygen species distribution: impact on Arabidopsis root system architecture. Mol Plant 8, 439–453.

Robinson, N.J., Procter, C.M., Connolly, E.L., and Guerinot, M.L. (1999). A ferric-chelate reductase for iron uptake from soils. Nature 397, 694–697.

Rodríguez-Celma, J., Connorton, J.M., Kruse, I., Green, R.T., Franceschetti, M., Chen, Y.T., Cui, Y., Ling, H.Q., Yeh, K.C., and Balk, J. (2019). Arabidopsis BRUTUS-LIKE E3 ligases negatively regulate iron uptake by targeting transcription factor FIT for recycling. Proc Natl Acad Sci U S A 116, 17584–17591.

Römheld, V., and Marschner, H. (1986). Evidence for a specific uptake system for iron phytosiderophores in roots of grasses. Plant Physiol 80, 175–180.

Roschzttardtz, H., Conéjéro, G., Curie, C., and Mari, S. (2009). Identification of the endodermal vacuole as the iron storage compartment in the Arabidopsis embryo. Plant Physiol 151, 1329–1338.

Roschzttardtz, H., Seguela-Arnaud, M., Briat, J.F., Vert, G., and Curie, C. (2011). The FRD3 citrate effluxer promotes iron nutrition between symplastically disconnected tissues throughout Arabidopsis development. Plant Cell 23, 2725–2737.

Santi, S., and Schmidt, W. (2009). Dissecting iron deficiency-induced proton extrusion in Arabidopsis roots. New Phytol 183, 1072–1084.

Scheepers, M., Spielmann, J., Boulanger, M., Carnol, M., Bosman, B., De Pauw, E., Goormaghtigh, E., Motte, P., and Hanikenne, M. (2020). Intertwined metal homeostasis, oxidative and biotic stress responses in the Arabidopsis frd3 mutant. Plant J.

Schmidt, H., Günther, C., Weber, M., Spörlein, C., Loscher, S., Böttcher, C., Schobert, R., and Clemens, S. (2014). Metabolome analysis of Arabidopsis thaliana roots identifies a key metabolic pathway for iron acquisition. PLoS One 9, e102444.

Schvartzman, M.S., Corso, M., Fataftah, N., Scheepers, M., Nouet, C., Bosman, B., Carnol, M., Motte, P., Verbruggen, N., and Hanikenne, M. (2018). Adaptation to high zinc depends on distinct mechanisms in metallicolous populations of Arabidopsis halleri. New Phytol 218, 269–282.

Selote, D., Samira, R., Matthiadis, A., Gillikin, J.W., and Long, T.A. (2015). Iron-binding E3 ligase mediates iron response in plants by targeting basic helix-loop-helix transcription factors. Plant Physiol 167, 273–286.

Shang, X., Cao, Y., and Ma, L. (2017). Alternative Splicing in Plant Genes: A Means of Regulating the Environmental Fitness of Plants. Int J Mol Sci 18.

Shanmugam, V., Lo, J.C., Wu, C.L., Wang, S.L., Lai, C.C., Connolly, E.L., Huang, J.L., and Yeh, K.C. (2011). Differential expression and regulation of iron-regulated metal transporters in Arabidopsis halleri and Arabidopsis thaliana - the role in zinc tolerance. New Phytol 190, 125–137.

Singh, S.K., and Reddy, V.R. (2017). Potassium Starvation Limits Soybean Growth More than the Photosynthetic Processes across CO2 Levels. Front Plant Sci 8, 991.

Sivitz, A.B., Hermand, V., Curie, C., and Vert, G. (2012). Arabidopsis bHLH100 and bHLH101 control iron homeostasis via a FIT-independent pathway. PLoS One 7, e44843.

Siwinska, J., Siatkowska, K., Olry, A., Grosjean, J., Hehn, A., Bourgaud, F., Meharg, A.A., Carey, M., Lojkowska, E., and Ihnatowicz, A. (2018). Scopoletin 8-hydroxylase: a novel enzyme involved in coumarin biosynthesis and iron-deficiency responses in Arabidopsis. J Exp Bot 69, 1735–1748.

Song, W.Y., Choi, K.S., Kim, D.Y., Geisler, M., Park, J., Vincenzetti, V., Schellenberg, M., Kim, S.H., Lim, Y.P., Noh, E.W., Lee, Y., and Martinoia, E. (2010). Arabidopsis PCR2 is a zinc exporter involved in both zinc extrusion and long-distance zinc transport. Plant Cell 22, 2237–2252.

Spielmann, J., Ahmadi, H., Scheepers, M., Weber, M., Nitsche, S., Carnol, M., Bosman, B., Kroymann, J., Motte, P., Clemens, S., and Hanikenne, M. (2020). The two copies of the zinc and cadmium ZIP6 transporter of Arabidopsis halleri have distinct effects on cadmium tolerance. Plant Cell Environ 43, 2143–2157.

Stacey, M.G., Patel, A., McClain, W.E., Mathieu, M., Remley, M., Rogers, E.E., Gassmann, W., Blevins, D.G., and Stacey, G. (2008). The Arabidopsis AtOPT3 protein functions in metal homeostasis and movement of iron to developing seeds. Plant Physiol 146, 589–601.

Talke, I.N., Hanikenne, M., and Krämer, U. (2006). Zinc-dependent global transcriptional control, transcriptional deregulation, and higher gene copy number for genes in metal homeostasis of the hyperaccumulator Arabidopsis halleri. Plant Physiol 142, 148–167.

Tehseen, M., Cairns, N., Sherson, S., and Cobbett, C.S. (2010). Metallochaperone-like genes in Arabidopsis thaliana. Metallomics 2, 556–564.

Thomine, S., and Vert, G. (2013). Iron transport in plants: better be safe than sorry. Curr Opin Plant Biol 16, 322–327.

Tsai, H.H., Rodríguez-Celma, J., Lan, P., Wu, Y.C., Vélez-Bermúdez, I.C., and Schmidt, W. (2018). Scopoletin 8-Hydroxylase-Mediated Fraxetin Production Is Crucial for Iron Mobilization. Plant Physiol 177, 194–207.

Verbruggen, N., and Hermans, C. (2013). Physiological and molecular responses to magnesium nutritional imbalance in plants. Plant and Soil 368, 87–99.

Vert, G., Grotz, N., Dédaldéchamp, F., Gaymard, F., Guerinot, M.L., Briat, J.F., and Curie, C. (2002). IRT1, an Arabidopsis transporter essential for iron uptake from the soil and for plant growth. Plant Cell 14, 1223–1233.

Wang, N., Cui, Y., Liu, Y., Fan, H., Du, J., Huang, Z., Yuan, Y., Wu, H., and Ling, H.Q. (2013). Requirement and functional redundancy of Ib subgroup bHLH proteins for iron deficiency responses and uptake in Arabidopsis thaliana. Mol Plant 6, 503–513.

Wintermans, J.F., and de Mots, A. (1965). Spectrophotometric characteristics of chlorophylls a and b and their pheophytins in ethanol. Biochim Biophys Acta 109, 448–453.

Xing, D., Wang, Y., Hamilton, M., Ben-Hur, A., and Reddy, A.S. (2015). Transcriptome-Wide Identification of RNA Targets of Arabidopsis SERINE/ARGININE-RICH45 Uncovers the Unexpected Roles of This RNA Binding Protein in RNA Processing. Plant Cell 27, 3294–3308.

Yi, Y., and Guerinot, M.L. (1996). Genetic evidence that induction of root Fe(III) chelate reductase activity is necessary for iron uptake under iron deficiency. Plant J 10, 835–844.

Yuan, Y., Wu, H., Wang, N., Li, J., Zhao, W., Du, J., Wang, D., and Ling, H.Q. (2008). FIT interacts with AtbHLH38 and AtbHLH39 in regulating iron uptake gene expression for iron homeostasis in Arabidopsis. Cell Res 18, 385–397.

Zhang, W., Du, B., Liu, D., and Qi, X. (2014). Splicing factor SR34b mutation reduces cadmium tolerance in Arabidopsis by regulating iron-regulated transporter 1 gene. Biochem Biophys Res Commun 455, 312–317.

Zhang, X.N., and Mount, S.M. (2009). Two alternatively spliced isoforms of the Arabidopsis SR45 protein have distinct roles during normal plant development. Plant Physiol 150, 1450–1458.

Zhang, X.N., Shi, Y., Powers, J.J., Gowda, N.B., Zhang, C., Ibrahim, H.M.M., Ball, H.B., Chen, S.L., Lu, H., and Mount, S.M. (2017). Transcriptome analyses reveal SR45 to be a neutral splicing regulator and a suppressor of innate immunity in Arabidopsis thaliana. BMC Genomics 18, 772.

